# Protein-primed RNA synthesis in SARS-CoVs and structural basis for inhibition by AT-527

**DOI:** 10.1101/2021.03.23.436564

**Authors:** Ashleigh Shannon, Véronique Fattorini, Bhawna Sama, Barbara Selisko, Mikael Feracci, Camille Falcou, Pierre Gauffre, Priscila El Kazzi, Etienne Decroly, Nadia Rabah, Karine Alvarez, Cécilia Eydoux, Jean-Claude Guillemot, Françoise Debart, Jean-Jacques Vasseur, Mathieu Noel, Adel Moussa, Steven Good, Kai Lin, Jean-Pierre Sommadossi, Yingxiao Zhu, Xiaodong Yan, Hui Shi, François Ferron, Bruno Canard

## Abstract

How viruses from the *Coronaviridae* family initiate viral RNA synthesis is unknown. Here we show that the SARS-CoV-1 and −2 **<u>*Ni*</u>**dovirus **<u>*R*</u>**dRp-**<u>*A*</u>**ssociated *N*ucleotidyltransferase (NiRAN) domain on nsp12 uridylates the viral cofactor nsp8, forming a UMP-Nsp8 covalent intermediate that subsequently primes RNA synthesis from a poly(A) template; a protein-priming mechanism reminiscent of *Picornaviridae* enzymes. In parallel, the RdRp active site of nsp12 synthesizes a pppGpU primer, which primes (-)ssRNA synthesis at the precise genome-poly(A) junction. The guanosine analogue 5’-triphosphate AT-9010 (prodrug: AT-527) tightly binds to the NiRAN and inhibits both nsp8-labeling and the initiation of RNA synthesis. A 2.98 Å resolution Cryo-EM structure of the SARS-CoV-2 nsp12-nsp7-(nsp8)_2_ /RNA/NTP quaternary complex shows AT-9010 simultaneously binds to both NiRAN and RdRp active site of nsp12, blocking their respective activities. AT-527 is currently in phase II clinical trials, and is a potent inhibitor of SARS-CoV-1 and −2, representing a promising drug for COVID-19 treatment.

## Introduction

Severe Acute Respiratory Syndrome coronavirus type 2 (SARS-CoV-2) is a human pathogen of the *Coronaviridae* (CoV) family, order *Nidovirales,* responsible for the ongoing pandemic which has so far resulted in >2.5 million fatalities (https://covid19.who.int/). This has prompted large-scale, global efforts which have greatly advanced our overall understanding of the virus and expedited vaccine development. However, much remains to be known regarding the specific mechanisms directing CoV replication and transcription of viral RNA, of direct importance for appropriate antiviral strategies.

Having ~30,000 nucleotides, the CoV positive-sense RNA (+RNA) genome is approximately three-times larger than that of significant human pathogenic +RNA viruses such as dengue, zika and poliovirus. This size difference reflects the acquisition of novel domains, many of which remain poorly characterized despite being potential candidates for CoV-specific drug targets. The viral genome is principally comprised of two-large open reading frames, Orf1a and Orf1ab, which are translated to yield 16 non-structural proteins (nsp1 – nsp16) responsible for viral replication and genome maintenance (Hartenian et al., 2020). Among these is the viral RNA-dependent-RNA polymerase (RdRp, nsp12), which associates with two small cofactor proteins, nsp7 and nsp8, to form the minimal replication-transcription complex (RTC) capable of RNA synthesis (Kirchdoerfer and Ward, 2019; Subissi et al., 2014). The 3’ end of the CoV genome specifies a nested set of subgenomic mRNAs which are translated to yield structural and accessory proteins.

Once expressed in the target cell, the viral RNA-dependent RNA-polymerase (RdRp) and associated proteins must diligently initiate viral RNA synthesis at precise termini to ensure all the genetic information is copied. RNA viruses have evolved a diverse range of initiation strategies, often dictated by subtle structural variations in the RdRp. Initiation is commonly divided into two main categories: *de novo* (primer-independent), and primer-dependent. However, specific mechanisms within these broadly defined groups can vary considerably. In the case of CoVs, the mechanism for initiation of RNA synthesis is poorly understood and somewhat controversial. The viral protein nsp8 is under scrutiny in this process, since it has been reported to synthesize short priming oligonucleotides (Imbert et al., 2006), carries an adenosine-specific terminal transferase (Tvarogová et al., 2019), and exhibits primer-extension activities (te Velthuis et al., 2012). On a structural level, the CoV RdRp nsp12 is related to “small-thumb” RNA polymerases, such as those of *Picornaviridae* (Peersen, 2019). These polymerases prime RNA synthesis through a ‘protein-primed’ mechanism (Paul et al., 1998), whereby a tyrosine hydroxyl group of a small viral peptide, known as VPg (Viral Protein genome-linked) is first covalently labeled with a uridine monophosphate (referred to as UMPylation). The VPg-pU is then extended to a dinucleotide and used to prime RNA synthesis, yielding genomic RNA which remains covalently linked to the viral VPg. Notably, the N-terminus of CoV nsp12 (just upstream of the RdRp domain) contains a Nidovirus-specific domain known as the NiRAN, which has been shown to mediate the covalent transfer of nucleoside monophosphates (NMP) to CoV nsp9 (Slanina et al., 2021). This activity was speculated to play a role in protein-priming, among other things (Lehmann et al., 2015), however so far there has been no evidence to support this, and the role of cofactor NMPylation in the viral life cycle remains unknown. More recently (and independent of its NMP-protein transferase activity), the NiRAN was proposed to participate in the guanylyltransferase step of viral RNA capping (Yan et al., 2021), suggesting it may play more than one role.

As with other RNA viruses, the universal set of core replicative enzymes, including the RdRp, remain at the forefront of drug-design strategies, with many clinically approved viral inhibitors currently being repurposed for attempted treatment of COVID infections. These include nucleoside analogues (NA). NAs play an important role in antiviral therapies because of their favorable drug properties: they can be administered as prodrugs which may be orally available and able to cross the blood-brain barrier. Once in the host cell, NA prodrugs are metabolised by cellular kinases into active 5’-triphosphate forms which compete with native nucleoside triphosphates (NTP) for incorporation into the viral RNA by the RdRp. Upon incorporation, NAs cause either chain-termination of RNA synthesis, or act as mutagenic nucleotides lethally altering the genetic make-up of the virus (Pruijssers and Denison, 2019). However, CoVs stand out amongst RNA viruses for possessing an RNA-repair 3’-to-5’ exonuclease (ExoN, nsp14) able to excise mismatched bases as well as chain terminating NAs (Bouvet et al., 2012; Eckerle et al., 2007, 2010; Ferron et al., 2018; Minskaia et al., 2006), generally compromising the efficacy of these drugs. While NAs are generally considered as RdRp-specific inhibitors, their potential to target other NTP-utilizing viral proteins, such as the NiRAN domain, has been largely unexplored.

Here, we investigate the role of the NiRAN domain of SARS-CoV in the initiation of viral RNA synthesis, and the potential for this domain to be targeted by NAs. We show that the CoV nsp12-nsp7-(nsp8)_2_ minimal RTC can initiate RNA synthesis through two distinct pathways: one protein-primed and mediated by the NiRAN domain through the UMPylation of nsp8, and the other through *de novo* synthesis of dinucleotide primers in a NiRAN-independent fashion. We additionally report the inhibition of both NiRAN transferase activity and *de novo* synthesis initiation with a promising guanosine NA candidate, AT-9010, the active triphosphate of the prodrug AT-527 (Good et al., 2021). Resolution of the nsp12-nsp7-(nsp8)_2_ Cryo-EM structure at 2.98 Å shows three target sites in the RTC: AT-9010 i) bound in the NiRAN active site; ii) incorporated at the +1 position of an RNA primer in the RdRp, and iii) poised for incorporation in the NTP binding site, confirming it is able to target several nucleotide binding sites of CoV-nsp12.

## Results

### The Nsp12 NiRAN domain UMPylates Nsp8

To elucidate the specifics behind SARS-CoV nsp12 nucleotidyl transferase activity, we incubated various combinations of SARS-CoV nsp12, nsp7 and nsp8, constituting the minimal RTC required for RNA synthesis (Subissi et al., 2014), in the presence of α^32^P-UTP. UMP is efficiently and specifically transferred to nsp8 in a reaction dependent on nsp12 and MnCl_2_ (Figure 1A, Figure S1A-C). Furthermore, a linked version of nsp7 and 8 (nsp7L8) is unable to be labeled by the NiRAN, indicated the authentic N-terminus of nsp8 is likely required. In contrast to the related equine arteritis virus (Lehmann et al., 2015), no covalent intermediate with nsp12 is formed. Various NiRAN mutants, including K73A, abolish labeling of nsp8 confirming that nucleotide transfer activity is provided by the NiRAN domain and not by nsp8 itself (Figure 1A,B, Figure S1D,E). The same specificity for nsp8-labeling was observed for the SARS-CoV-2 RTC (Figure S2F). A comparison of nsp8 labeling efficiency with the four native nucleotides shows UTP is the preferred substrate, albeit with a structural flexibility which allows the binding and transfer of GTP, CTP and ATP to a lesser extent (Figure 1C). To determine the labeling site, stability of the UMP-nsp8 bond was assessed at high or low pH (Figure 1D). The UMP-nsp8 bond is stable in 0.1M HCL, but alkali labile, indicating that most of the UMP is bound to the hydroxyl group of either a serine or threonine residue. Interestingly however, when the nsp8 labeling reaction is performed in the presence of a poly(A)_27_ oligoribonucleotide and nsp7 (i.e. the nsp12-nsp7-(nsp8)_2_ RTC), the UMP-nsp8 bond is stable at both high and low pH, indicative of a phosphodiester bond with a tyrosine hydroxyl group (Figure 1D). Only two serines (S11 and S85) and one tyrosine (Y71) are highly conserved among CoV nsp8 (Subissi et al., 2014). Single and double alanine-mutants of these residues do not eliminate labeling; Notably, we detect an increase in labeling efficiency for S11A suggesting that this residue may participate in the selectivity of the labeling site (Figure 1B). This data suggests that several residues of nsp8 can be labeled by the NiRAN, and that this is depended on the conformation of the RTC, dictated by the presence or absence of RNA. Recent Cryo-EM structure models show nsp8 forming ‘molecular sliding poles’ stabilizing the RNA exiting from the polymerase active site (Hillen et al., 2020). In the absence of RNA, these flexible N-terminal extensions of nsp8 would be in an alternative conformation, altering the nucleotidylation site, and potentially serving as a mechanism to regulate NiRAN activity.

**Figure 1.**
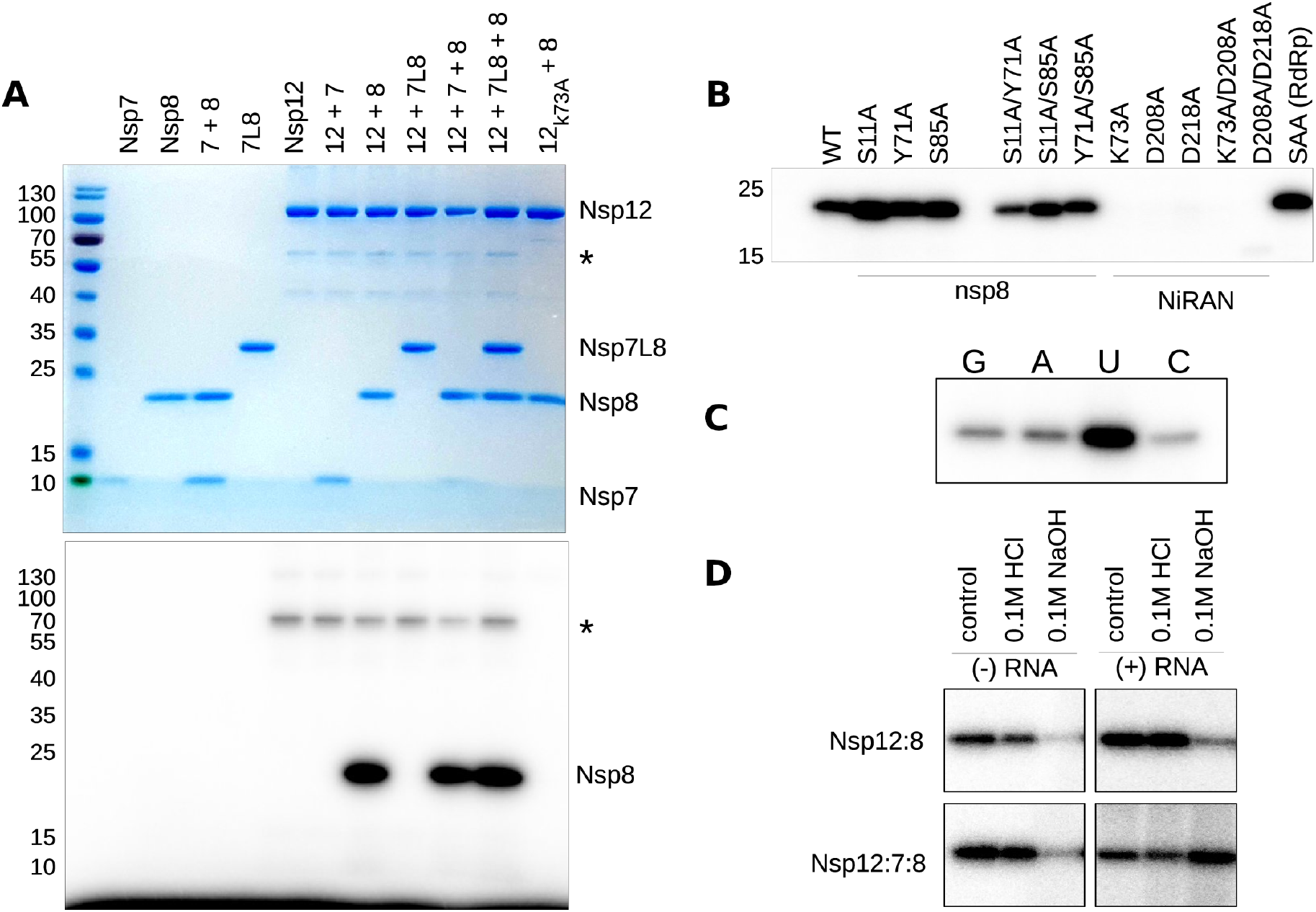
The nsp12 NiRAN domain mediates transfer of NMPs to viral cofactor nsp8. (A) Various combinations of nsp12, nsp7 and nsp8, as well as a covalently linked version of nsp7 and 8 (nsp7L8) were incubated for 1hr at 37°C with α^32^P-UTP. Samples were separated on a 15% SDS PAGE gel to remove non-covalently bound nucleotides and stained for total protein (top), then exposed to reveal radioactivity (bottom). Nsp8 and a small amount of contaminating protein (*) is labeled in a nsp12-dependent fashion. A NiRAN active site mutant (K73A) eliminates all labeling (rightmost lane). (B) Labeling of various nsp8 mutants by nsp12, and of nsp8 WT by various nsp12 NiRAN mutants. Final lane shows RdRp SAA active site mutant. (C) Comparison of nsp8-labeling efficiency by nsp12 with four α^32^P-radiolabelled nucleotides. (D) Stability of the nsp8-UMP bond, assessed through treatment at low or high pH. The labeling reaction was performed with either nsp12 + 8 alone (top gels) or with the RTC (bottom gels) in the absence (left) or presence (right) of poly(A)_27_ RNA.

### SARS-CoV UMPylated nsp8 is involved in a unique protein-primed RNA synthesis step

We recently established a high-throughput, fluorescent screening assay using the purified SARS-CoV RTC, able to synthesise poly(U) from a poly(A) template in the absence of a primer (Eydoux et al., 2021). To gauge how synthesis is initiated in these assays, and whether the UMPylation of nsp8 is involved, we incubated the nsp12-nsp7-(nsp8)_2_ RTC with poly(A)_27_ oligo-ribonucleotide and UTP (supplemented with α^32^P-UTP), and separated the products on denaturing SDS-PAGE and high-resolution urea PAGE sequencing gels (Figure 2A,B and Figure S2A). The RTC efficiently synthesizes labeled RNA in the presence of MnCl_2_ (Figure S1B,C), with the majority of products longer than the expected Poly(U)_27_ size (Figure 2B, Figure S2A). Modification of either the 5’ or 3’ end of the poly(A)_27_ template does not affect synthesis, demonstrating that these products are not a result of nsp8 terminal-transferase activity (Tvarogová et al., 2019), or other template modification (Figure S2B). Rather, once a full-length poly(U)_27_ product is synthesized, the complex is able to switch to a new poly(A)_27_ acceptor template and continue synthesis, resulting in products that are multimeric in length to the input poly(A)_27_ template. Similar template-switching activity has been previously described for other viral RdRps (Menéndez-Arias et al., 2017; Woodman et al., 2016). Despite the previously reported primase and polymerase-related activities of nsp8 (Imbert et al., 2006; Tvarogová et al., 2019; te Velthuis et al., 2012), we do not observe activity of nsp8 alone or in combination with nsp7 in this context (Figure 2A, Figure S2A). We additionally noted the presence of radiolabeled product either in the wells, or with very limited migration down the gel (Figure 2B, red arrow). Proteinase K (PK) hydrolysis of the polymerase complexes lead to the digestion of these products and results in the further release of poly(U) synthesized RNA, consistent with an nsp8-primed synthesis event. Western blot analysis with anti-nsp8 confirms that the higher products (>40 kDa on SDS PAGE) are covalently linked with nsp8, and not another protein (Figure S2C, lanes 1-2). Furthermore, the specificity of the labeling was demonstrated by mutating the K73A of the NiRAN, showing this reaction is NiRAN-dependent (Figure S2C, lane 11). Interestingly, polymerization can also be initiated on poly(U) and poly(C) templates through the addition of ATP and GTP, respectively (Figure S2D,E). However, in contrast to poly(U) synthesis, poly(A) and poly(G) products are not retained in the wells, and furthermore are not sensitive to proteinase K digestion. Thus, protein-primed RNA synthesis is UTP-specific, consistent with the preference for UMP-nsp8 labeling by the NiRAN domain. We conclude that the RTC complex UMPylates one of its presumably bound nsp8, which serves as an uridylated primer to initiate poly(U)_n_ synthesis.

**Figure 2.**
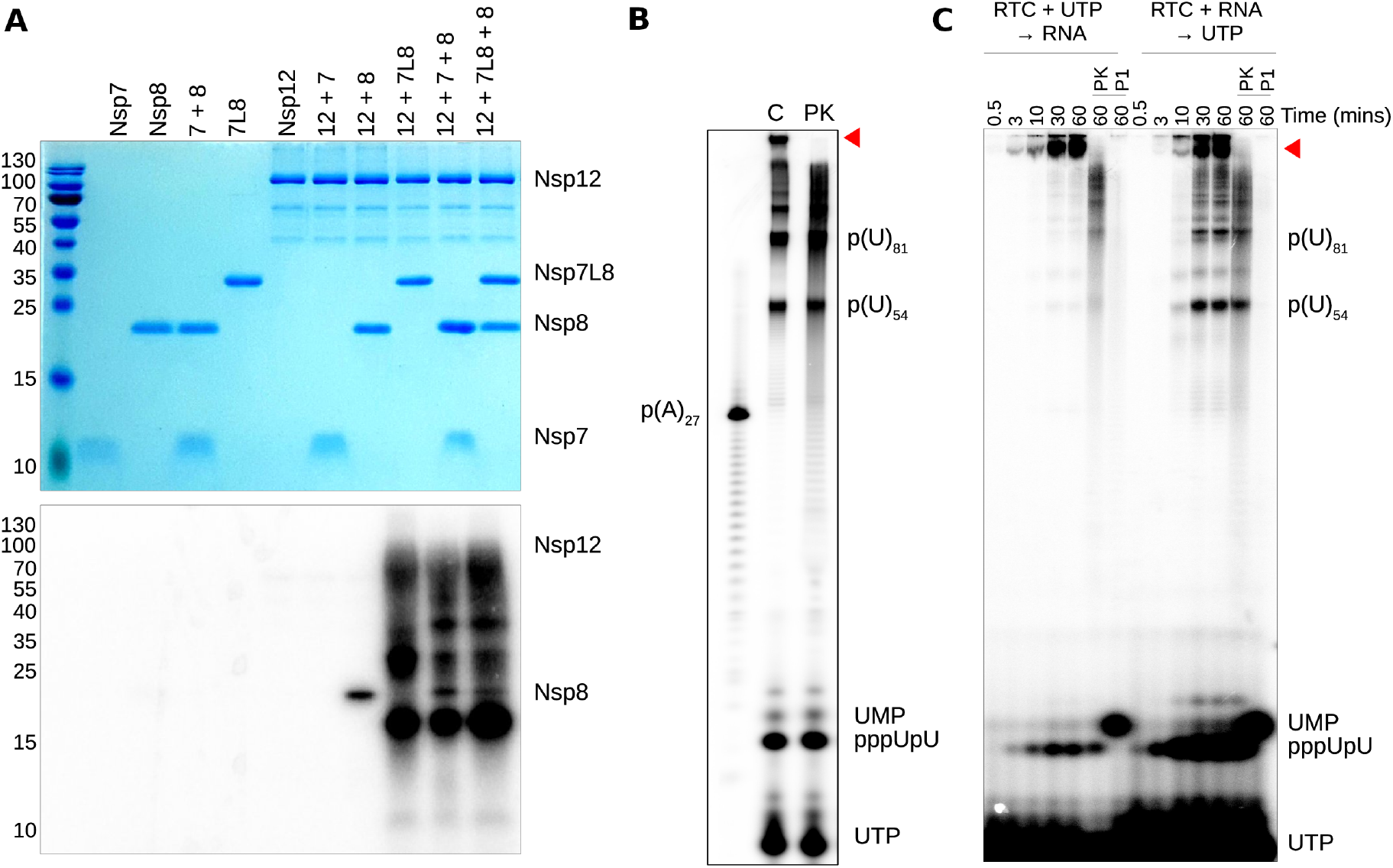
The RTC initiates primer-independent synthesis of long poly(U)_n_ RNA from a poly(A)_27_ template. (A) Various combinations of nsp12, nsp7 and nsp8, and nsp7L8 were incubated for 1hr at 37°C with UTP (supplemented with α^32^P-UTP) and poly(A)_27_ RNA. Samples were separated on a 15% SDS PAGE gel to remove non-covalently bound nucleotides/RNA and stained for total protein (top), then exposed to reveal radioactivity (bottom). Equivalent urea-PAGE shown in Figure S2A. (B) Activity of the full RTC, as described in (A), separated on a 14% acrylamide 7M urea gel. Size of input template RNA shown as p(A)27, with p(U)_54_ and p(U)81 showing multimeric poly(U) synthesis products. C represents reaction control, while PK shows same sample following protein digestion with proteinase K. (C) Order of addition experiment. The nsp12:7:8 RTC was incubated for 30 min at 37 °C with either UTP or RNA. Following incubation, the complementary reagent was added and reactions were stopped at indicated time-points. Reactions run for 60 min were additionally treated with proteinase K (PK) or nuclease P1 (P1) for protein and RNA digestion, respectively.

### Two independent pathways of primer synthesis co-exist to initiate RNA synthesis

For the *Picornaviridae* family, the VPg protein is used to prime both plus and minus strand synthesis. Its covalent attachment to the 5’ end of the RNA additionally substitutes for the RNA-cap, protecting the viral RNA from host cell degradation. In contrast, the CoV genome presumably contains a conventional ^m7^GpppA_2’Om_ RNA cap-1 structure (Bouvet et al., 2010). This difference suggests that for CoVs, the nsp8-UMP protein-primed strategy is specific to minus strand synthesis, templated from the poly(A) tail. We noted that NiRAN mutants that completely eliminate nsp8 UMPylation are still able to synthesise poly(U) RNA, although notably with a loss in high-molecular weight products (Figure S3A). To address this, we set-up an order-of-addition experiment, incubating the RTC either with UTP or poly(A)_27_ RNA first, prior to addition of the complementary reagent. Pre-incubation with UTP, promoting NiRAN-mediated nsp8-UMPylation, yields exclusively high-molecular weight poly(U)_n_ products covalently bound to nsp8 (Figure 2C, red arrow, Figure S3B). In contrast, pre-incubation of the RTC with poly(A)_27_ results in production of both protein-bound RNA (released by PK digestion), and long poly(U)_n_ products not bound to nsp8 (Figure 2C). NiRAN mutants which eliminate UMP-nsp8 labeling also eliminate higher molecular-weight protein-primed products, but do not abolish the synthesis of poly(U) products (Figure S3A,C-F). This indicates that two distinct priming mechanisms coexist: one NiRAN dependent promoted by UMP-nsp8 (pathway 1), and the other NiRAN independent (pathway 2).

### The SARS-CoV RTC synthesizes 5’-triphophate dinucleotide primers in a NiRAN-independent manner

We noted that for NiRAN mutant RTCs, polymerization through pathway 2 was actually increased, indicating that the two initiation mechanisms occur simultaneously and in competition (Figure S3C). NiRAN mutant RTCs also produced higher quantities of a small molecular-weight UMP-containing product, thus potentially involved in the second priming pathway. This product was identified as pppUpU based on its sensitivity to Calf Alkaline phosphatase and nuclease P1, as well as co-migration with chemically synthesized pppUpU. Addition of free pppUpU to the RTC increases synthesis of unbound poly(U) n RNA products, and decreases the lag time of the reaction in a concentration dependent manner, showing that synthesis of a dinucleotide pppUpU is a prerequisite for pathway 2 (Figure S4A,B). As expected, an RdRp active site mutant of nsp12 (SDD -> SAA) abolishes poly(U) synthesis, and additionally eliminates synthesis of pppUpU (Figure S3C). This confirms that it is the RdRp domain of nsp12, in conjunction with nsp7 and 8, that drives production of this dinucleotide primer, and not the NiRAN.

### Nsp12-mediated pppGpU synthesis directs the precise start of primer elongation on (-)ssRNA sequence

To better understand how the two pathways are regulated during minus strand synthesis, we used a heteropolymeric RNA corresponding to the last 20 nucleotides of the 3’-end of the genome (ST20) with or without a poly(A)_15_ tail. In the absence of the poly(A) sequence, spurious self-primed products are synthesized, and can be eliminated by blocking the 3’ end of the template (Figure S4C). Strikingly, we observe that in the presence of the poly(A)_15_ tail, synthesis is initiated in the vicinity of the RNA-poly(A) junction, rather than from the end of the poly(A) sequence as judged by the size of the synthesis product (Figure 3A, -ve panel). This indicates that the poly(A) tail and 3’ genomic RNA sequence elements guide the positioning of the RTC to the true 3’-end of the genome, i.e., at its junction with the poly(A) tail, for initiation of synthesis. Pre-incubation of the complex with UTP can additionally promote low-level synthesis of the full-length template (ST20 + poly(A)_15_) through the protein-primed pathway 1, which is released following proteinase K digestion (Figure S4C). Western blot analysis confirms that the full-length RNA product is covalently attached to nsp8, and that this is dependent on the NiRAN domain (Figure S2C).

**Figure 3.**
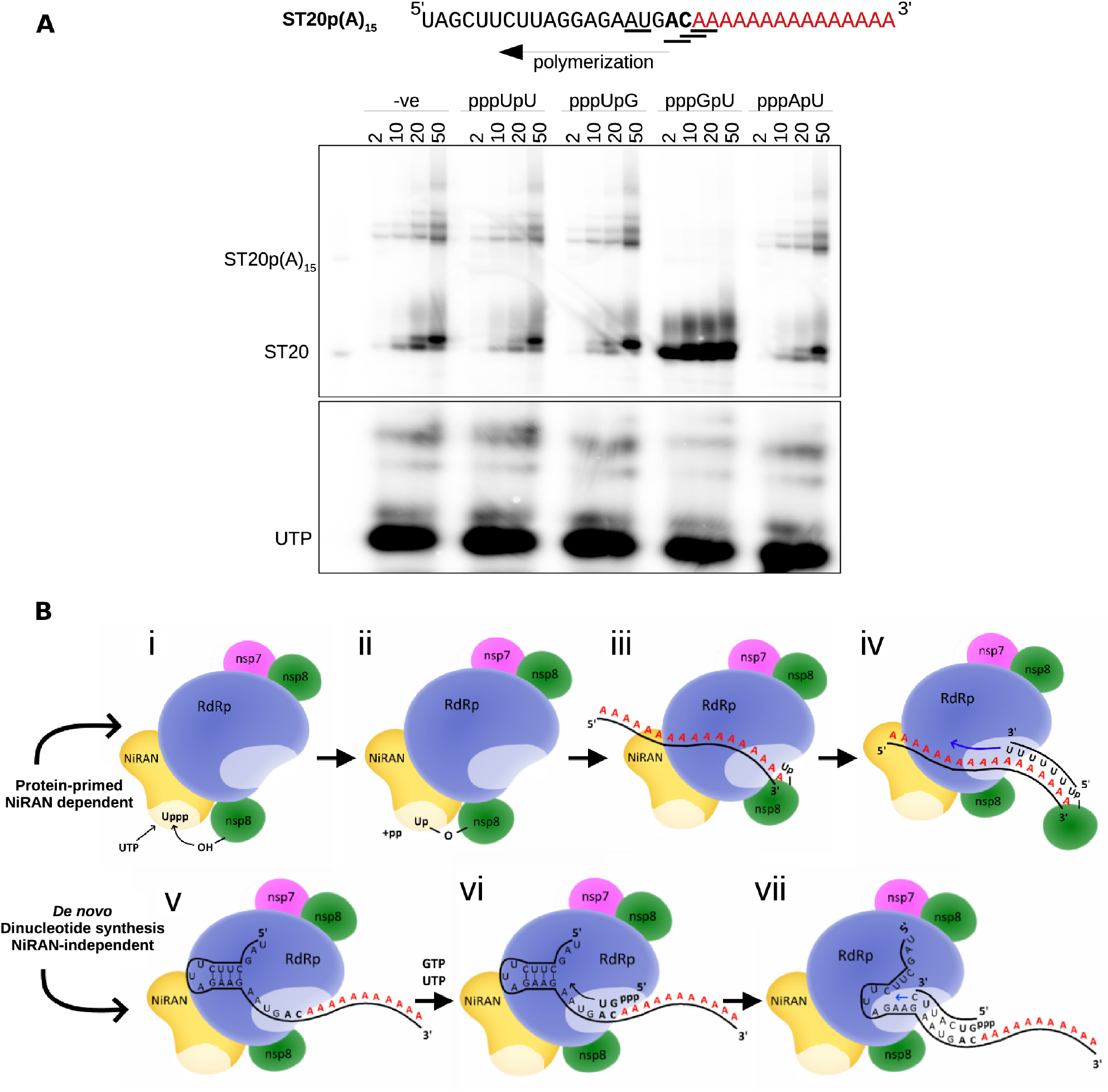
RNA synthesis is preferentially directed to the genome-poly(A) junction sequence and shows preference for pppGpU. (A) An RNA template mimicking the SARS-CoV-1 or 2 genome-poly(A) junction sequence was incubated with the RTC, NTPs (supplemented with α^32^P-UTP), and 100 μM of chemically synthesized, dinucleoside 5’-triphosphates as indicated. Control in the absence of a dinucleotide shown as (− ve). Synthesis products were separated using 7M urea-PAGE (20 % acrylamide) and visualized using a Typhoon FluorImager. (B) Two RNA synthesis initiation pathways by the SARS-CoV RTC. In pathway 1, NiRAN-bound UTP (i) is transferred to nsp8 to yield UMP-nsp8 (ii), further positioned to the poly(A) 3’-end (iii) to prime RNA synthesis (iv). In pathway 2, the ST20p(A)_15_ sequence drives nsp12 RdRp active site binding at the heteropolymeric – polyA junction (v) to synthesize a pppGpU dinucleotide primer (vi) able to prime RNA synthesis of the complementary strand (vii). ST20p(A)_15_ secondary structure based on RNAfold WebServer prediction.

To determine the precise sequence initiation site, we supplemented the reaction with various chemically synthesized pppNpN dinucleotide primers. Addition of pppGpU, the sequence complementary to the last two bases immediately adjacent to the poly(A) tail, greatly increased both the reaction rate and level of product formation (Figure 3A). In contrast, pppUpU, pppUpG and pppApU dinucleotides had a minimal effect on the reaction. We conclude that the polymerase complex preferably initiates synthesis with a pppGpU dinucleotide, templated from the precise 3’ end of the genome-poly(A) junction, immediately upstream the poly(A) tail. Synthesis of this dinucleotide is presumed to be the rate-limiting step, and can be surpassed through the addition of free pppGpU. Interestingly, in contrast to pppUpU, free pppGpU is not readily observed, indicating that it remains bound in the polymerase active site and is rapidly extended upon production. In contrast, pppUpU appears to be synthesized more efficiently, however it is released by the polymerase RTC, indicating that it is sub-optimal to start (-)RNA synthesis. Figure 3B recapitulates both pathways operating in parallel at the 3’-end of the SARS-CoV genome. Altogether, our results show that the SARS-CoV RTC promotes RNA synthesis initiation through two distinct pathways: pathway 1 is through protein-priming reminiscent of *Picornaviridae*, and pathway 2 is by means of *de novo* synthesis of pppNpN primers, pppGpU being preferred to start at the genome-poly(A) junction.

### AT-9010 and STP terminate RNA synthesis but are excised by SARS-CoV nsp14 Exonuclease

Given the specific role of UTP and GTP in the (-)RNA priming event, we sought to determine whether uracil- or guanine- nucleoside/tide analogues (NAs) could inhibit priming activity. Since most NAs generally exert an antiviral effect by targeting the viral RdRp for incorporation into viral RNA (Pruijssers and Denison, 2019), it was of particular interest to establish whether the NiRAN domain would additionally constitute an antiviral target.

We investigated the inhibition potential of the active metabolites of two NAs; the uracil analogue Sofosbuvir (SOF) and its guanosine equivalent AT-511. SOF is clinically approved for the treatment of hepatitis C virus (HCV) (Dousson, 2018), however has shown limited efficacy against SARS-CoV-2 infection (Good et al., 2021; Han et al., 2021), Han 2021). In contrast, AT-527 (the hemi-sulfate salt of AT-511) was recently shown to act as a potent broad-spectrum anti-CoV inhibitor in a variety of cell lines. It is now in phase II clinical trials for the treatment of both HCV infection (Good et al., 2020) and COVID-19 (Good et al., 2020).

Both SOF and AT-511 are phosphoramidate prodrugs containing a 2’-fluoro-2’-C-methyl modified ribose, with the only difference being the nucleobase (Figure S5A). In cells, these prodrugs are metabolized by cellular kinases into active 5’-triphosphate forms (AT-9010 and STP, respectively), which presumably act as substrates for the viral RdRp for incorporation into viral RNA, as has been shown for HCV (reviewed in (Dousson, 2018)). We first compared the RdRp selectivity for these two NAs using the SARS-CoV RTC and a heteropolymeric RNA primer/template pair (Shannon et al., 2020). Both AT-9010 and STP are incorporated into RNA by the RdRp as a substitute for GTP or UTP, respectively, causing immediate chain termination (Figure S5B). In the presence of GTP, AT-9010 is a competitive guanosine substrate, discriminated 22-fold against GTP (Figure S5C). In contrast, STP:UTP competition experiments (20:1) show STP is not competitive at this ratio (Figure S5D). However, following incorporation into RNA, both drugs are excised by the SARS-CoV ExoN (Figure S5D), indicating that the SARS-CoV RTC proofreading activity potentially jeopardizes their efficacy. It therefore seems unlikely that the potent anti-CoV activity of AT-527 would solely be provided by RdRp-mediated incorporation of AT-9010 into RNA.

### AT-9010 binds to the NiRAN active site, inhibiting nsp8-UMPylation (Pathway 1)

To determine whether either drug was able to additionally target the NiRAN transfer activity, we performed competition experiments measuring the efficiency of nsp8-UMPylation in the presence of increasing concentrations of AT-9010 or STP (Figure 4A, Figure S6A-C). Both drugs were able to inhibit labeling, in contrast to ^m7^GTP, used as a control. Interestingly, despite the preferential labeling of nsp8 with UMP, the uracil analogue STP is ~4-5-fold less efficient at blocking nsp8 labeling than AT-9010, consistent at two enzyme concentrations (1 μM and 5 μM nsp12, with 5-fold excess nsp8) (Figure 4A, Figure S6B). The calculated IC_50_ (half maximal inhibitory concentration) of inhibition of nsp8-UMPylation by AT-9010 gives IC_50_ values of approximately half the concentration of nsp12 (0.87 and 1.9 μM, respectively), indicating that AT-9010 binds to the NiRAN domain at a roughly 1:1 stoichiometry, strongly outcompeting UTP. Furthermore, given the excess of nsp8 in the reaction, these results suggest that AT-9010 remains stably bound in the NiRAN active site, rather than being transferred to nsp8. Both STP and AT-9010 were additionally shown to inhibit GMPylation of nsp8 (Figure S6C).

**Figure 4.**
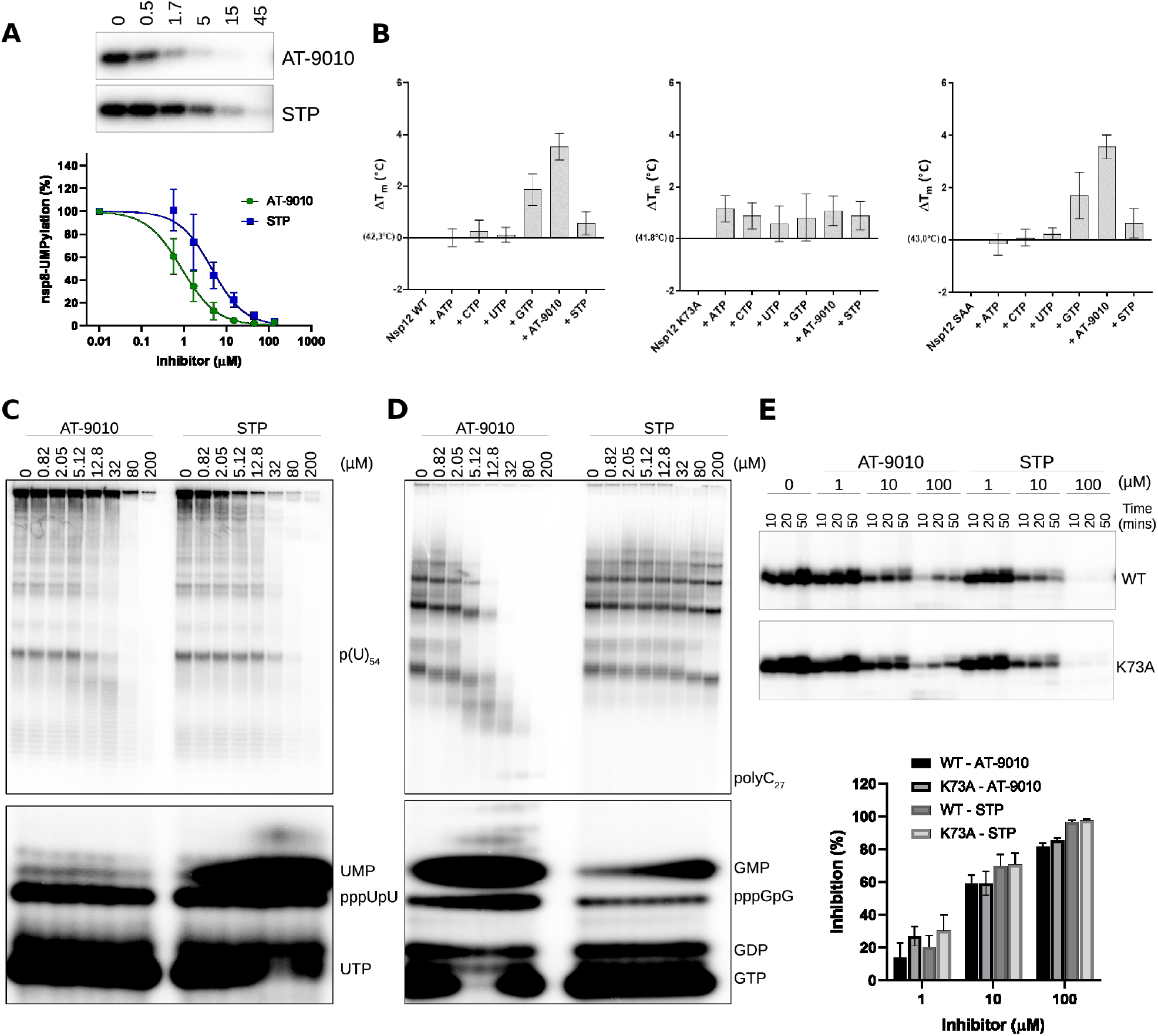
AT-9010 inhibition of both protein-primed and *de novo* synthesis. (A) Labeling of nsp8 by nsp12 with a 5 μM constant concentration of UTP (supplemented with α^32^P-NTP), in competition with increasing concentrations of AT-9010 or STP (n=3, SD shown). Gel shows representation of 3 individual data sets. Total intensity was quantified with ImageQuant, and plotted at % residual activity. Calculated IC 50 values using 1 μM nsp12 with 5-fold molar excess of nsp8 were 0.87 ± 0.1 for AT-9010 and 4.5 ± 0.7 for STP (5-fold difference). (B) Thermal stability of nsp12 WT (left), NiRAN mutant K73A (middle) and RdRp mutant SAA (right) in the presence of different native NTPs or inhibitors. Reactions were run with 5 mM MgCl_2_, and 0.5 mM MnCl_2_ in triplicate, SD shown. (C-D) Inhibition of nsp12:7:8 RTC synthesis from poly(A)_27_ (c) and poly(C)_27_ (d) templates with AT-9010 and STP. Gel is representation of 2 independent experiments. (E) Inhibition of synthesis from ST20p(A)_15_ template by RTC containing either WT or NiRAN mutant (K73A) nsp12. Graph represents average activity from two independent experiments, with calculated difference from three time-points compared to no-inhibitor control. Band shown corresponds to ST20 product with synthesis starting at the poly(A) junction.

Thermal shift assays with nsp12 in the presence of MnCl_2_ confirms that AT-9010 provides more thermodynamic stability than any other native nucleotide (Figure 4B). Comparison of NiRAN and RdRp active site mutants (K73A and SAA, respectively) shows that this stability increase is provided by AT-9010 binding preferentially into the NiRAN active site, rather than the RdRp active site. Both GTP- and AT-9010-nsp12 complexes show an increase in stability compared with UTP or STP-bound complexes. Consistent with inhibition results, these results indicate that guanosine is the preferred base of the NiRAN active site, and the 2’-fluoro-2’-C-methyl ribose modification of AT-9010 provides additional stability.

### AT-9010 inhibits RNA synthesis through pathway 2

We additionally tested the ability of the two drugs to inhibit initiation of RNA synthesis from a poly(A)_27_ template. Despite AT-9010 being a guanosine analogue, it inhibits synthesis of poly(U)_n_ RNA equally as well as STP (Figure 4C). The inhibition by AT-9010 shows that the majority of its inhibitory activity might not be due to RdRp-mediated incorporation, as purine-purine mismatches (in this case AT-9010:A, equivalent to G:A) would be significantly disfavored over the native UTP:A. Inhibition of RNA synthesis is comparable with the nsp12 K73A mutant, indicating that inhibition cannot be allosteric via binding to the NiRAN domain (Figure S6D). Rather, inhibition occurs through blocking RNA synthesis initiation at the RdRp active site. The equivalent experiment with a poly(C) template, theoretically favoring AT-9010, shows STP to be virtually inactive, while AT-9010 significantly reduces activity (Figure 4D). It therefore appears that while both drugs are able to inhibit *de novo,* NiRAN-independent initiation of RNA synthesis, AT-9010 inhibition is somewhat template-independent. Preincubation of the complexes with either STP or AT-9010, prior to addition of pppGpU and other NTPs inhibits synthesis of ST20p(A)_15_ RNA at similar levels. Again, inhibition is comparable with WT and K73A NiRAN mutant complexes, showing that both drugs are additionally able to bind in the RdRp active site and prevent synthesis and/or binding of dinucleotide primers (Figure 4E).

### Structural basis for nsp12 inhibition by AT-9010

Previously reported structures of the SARS-CoV-2 RTC in the presence and absence of RNA (Gao et al., 2020; Wang et al., 2020) have paved the way to understand inhibition at a structural level (Yin et al., 2020). To investigate the binding and incorporation into RNA of AT-9010, and in particular the relationship between the RdRp and NiRAN active sites in its presence, we performed Cryo-EM studies using a dsRNA-bound to the SARS-CoV-2 RTC in the presence of AT-9010. Image processing of single particles allowed the reconstruction of a RTC/RNA/AT-9010 quaternary complex at 2.98 Å resolution (Table S1, Figure S7), showing AT-9010 simultaneously binding to both NiRAN and RdRp active site of nsp12 (Figure 5). The overall structure resembles previously reported structures, with one nsp12, one nsp7 and two nsp8 proteins (Figure 5A). The RdRp domain is bound to a primer-template dsRNA pair, with the 5’-monophosphate of AT-9010 incorporated at the 3’ end of the RNA primer (Figure 5B,C). A second, free AT-9010 triphosphate is present, being loaded into the polymerase at the NTP binding site (framed from left by motif C, bottom by motifs A, D, and top by motif F) (Figure 5C,D). Residues Asp618 and Asp760 of motifs A and C, respectively, coordinate a single catalytic magnesium ion, which interacts with the phosphates of the second AT-9010 (Figure 5D, Figure 6). Finally, the structure also shows AT-9010, bound in the active site of the NiRAN domain (Figure 5E).

**Figure 5.**
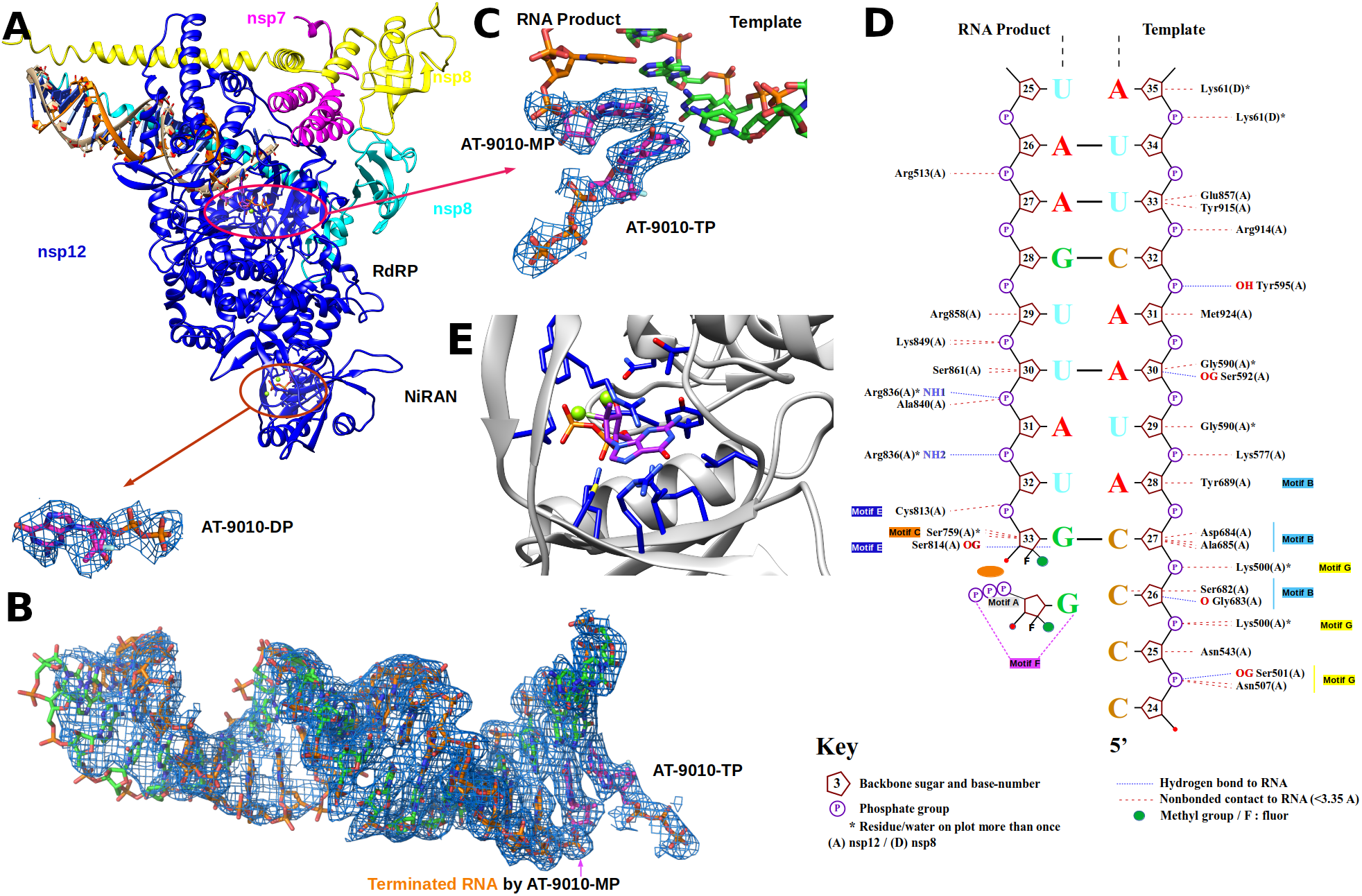
Cryo-EM structure of the RTC with bound RNA and AT-9010 molecules. (A) Ribbon representation of the Cryo-EM structure of nsp7-(nsp8)_2_-nsp12:AT-9010-terminated-RNA:(AT9010)2 complex. Red ovals are enlarged in (B) and (C) with density map and stick representation of two AT9010 molecules in the RdRp active site. One AT-9010 is covalently incorporated into the RNA strand, (D) Nucplot molecular analysis between RNA AT-9010 and nsp12 (E) AT-9010 in its diphosphate form (sticks) bound to the NiRAN domain. Ribbons are depicted as follows: nsp12: blue; nsp7: magenta; nsp81 and nsp82: yellow and cyan, respectively; RNA: green sticks; AT9010: magenta sticks.

**Figure 6.**
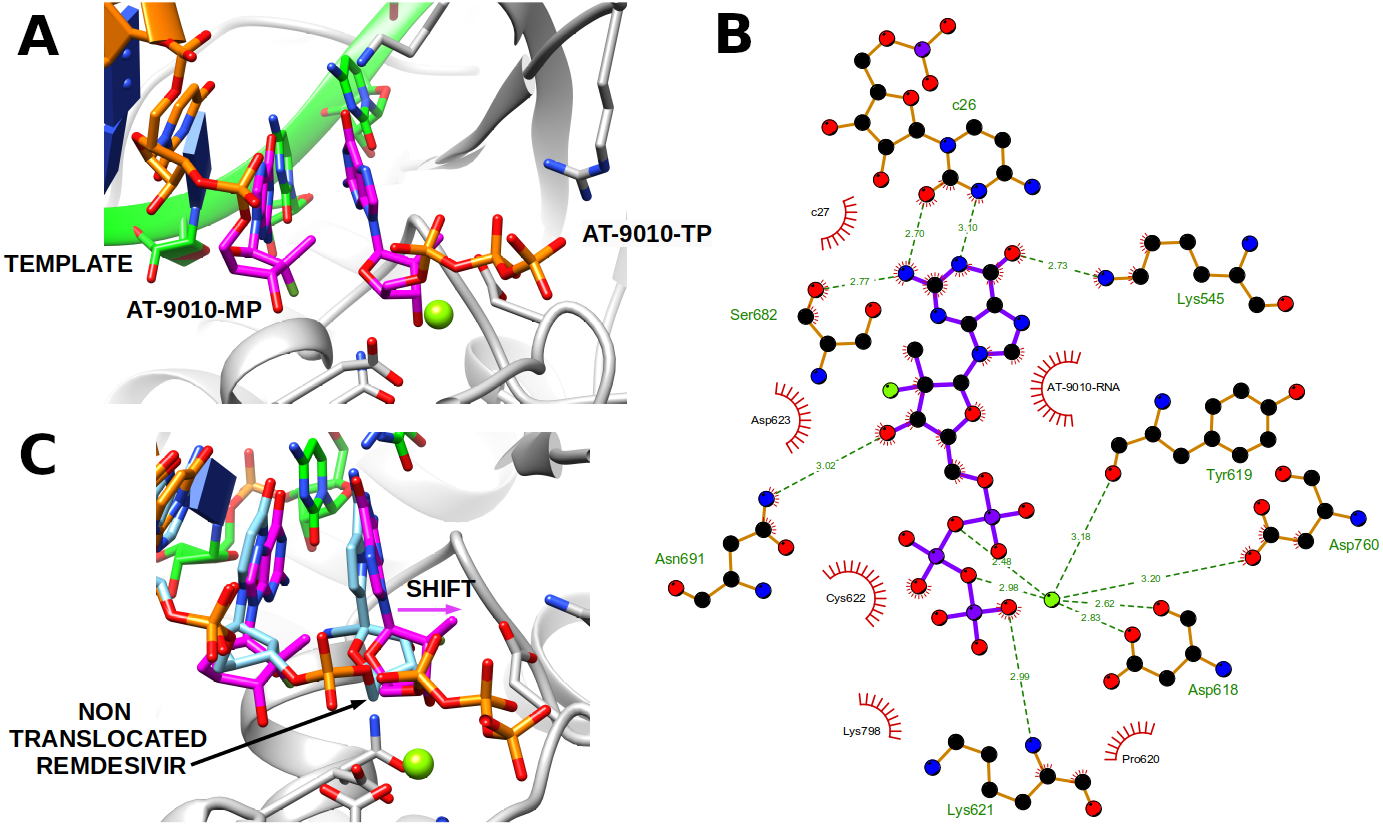
Structural basis for inhibition of the SARS-CoV RTC by AT-9010. (A) One AT-9010 5’-monophosphate is incorporated into the primer RNA strand, and terminates RNA elongation. The second AT-9010 molecule coordinated by one ion is shown in the NTP binding site in stick representation. (B) Ligplot 2D analysis of the contacts of AT-9010 molecules. (C) Superimposition of AT-9010 molecule with a terminally incorporated, untranslocated NMP, here remdesivir (PDB: 7BV2). The position of the ribose is pushed away and phosphates are in a post-incorporation position.

### The nsp12 active site incorporates AT-9010 into RNA and terminates synthesis

The visible 5’-end of the RNA templates is composed of 4 consecutive cytidine bases (C24-C27) designed to favor the incorporation of AT-9010 into the RNA product strand. AT-9010 is incorporated at position +1 of the RNA product, pairing with C27 of the template strand. A second AT-9010 is observed in a pre-incorporation state partially pairing with C26 of the template, and representing the first snapshot of a SARS-CoV-2 polymerase inhibited as such: the RdRp-RNA complex is stalled in a post-translocation state, with the −1 site is occupied by the second AT-9010, poised for incorporation, and preventing other NTPs from being loaded (Figure 5D, Figure 6A).

The guanine of the incorporated AT-9010 is canonically base-paired with the cytosine (C27) of the template (Figure 5C,D). The ribose forms hydrogen bonds with Ser814 and the 5’ phosphate, coordinated by Cys813. The effect of the 2’-fluoro-2’-C-methyl ribose modification is two-fold. Firstly, the replacement of the 2’ hydroxyl by a fluoro group eliminates the interaction between Ser759 (motif C) and the 2’-OH, which usually stabilizes the ribose. This sterically displaces the catalytic ions that are normally coordinated by residues of motif C and the RNA product. This observation is corroborated by the lack of a second catalytic Mg^2+^ in the structure, that is usually coordinated by the catalytic Asp760 in motif C and with the 3’-OH and phosphate of the last incorporated nucleotide of the RNA product (Figure 6B). This ion plays a critical role in the positioning of the 3’ end of the RNA product and the incoming NTP. Secondly, the hydrophobic methyl group of the AT-9010 ribose amplifies the inhibitory effect by creating a hydrophobic hindrance that prevents correct positioning of the ribose of the incoming NTP. This prevents further elongation of the product strand, irrespective of the presence of a 3’-OH, explaining why AT-9010 acts as a chain terminator. As a result, the second triphosphate AT-9010 is stalled at the −1 site position. The ribose ring is pushed away compared to its theoretical position due to hydrophobic repulsion, mostly driven by the methyl group of the incorporated AT-9010 (Figure 6C). As a result, the side chain of Asp623 (motif A), usually responsible for stabilising the pyrophosphate after incorporation, is also pushed away. The guanine is partially base-paired with the cytosine (C26) of the template strand, and is further stabilized by residues Lys545 (motif F) and Ser682 (motif B), two residues important during the fidelity check, prior to incorporation (Figure 5D, Figure 6B).

The α- and β- phosphates are coordinated by the Mg^2+^ and the γ-phosphate is stabilized by the Lys621 also in motif A and Lys798 in motif D. The distances between the α-phosphate and the 3’-OH necessary to allow metal coordination and the bonding event (for incorporation) are not respected, preventing incorporation. Rather we observe that AT-9010 has its α- and β -phosphates spatially overlapping the position of a pyrophosphate (β- and γ-phosphates) (Figure 6C).

### The NiRAN domain binds AT-9010 at the UMPylation active site

The NiRAN domain is structurally related to the pseudo-kinase family of enzymes (Slanina et al.,2021), allowing delineation of the catalytic residues of the enzyme (Figure S8A).However,CoV-unique sequence and structural elements can also be identified for the NiRAN domain. The structure shows AT-9010 in its diphosphate form, bound to the NiRAN domain. The binding site is made of a closed cavity,which opens into a groove harbouring two catalytic ions coordinated by the conserved Asn209 and Asp218.The groove further widens to a flat surface formed by the 2 β-strands (β2-β3 residues 33-48) (Figure S8B,C). The base and the modified ribose of AT-9010 fits snuggly in the cavity (Figure 7A,B), while the α and β phosphates are coordinated in the groove by the two catalytic ions and Lys73. The guanine base is intensively stabilized by hydrophobic interaction and through hydrogen bonding with residues Arg55, Thr120 and Tyr217, residues which are conserved in CoV- NiRAN sequences (Figure 7), but are not present in other pseudo-kinases (Figure S8A). The binding mode of AT-9010 is reminiscent of the orientation of ATP bound to the casein kinase (Xu et al., 1995) but strikingly different from the position of the nucleotide in the pseudo-kinase structures (Sreelatha et al., 2018; Yang et al., 2020) and a recently published GDP-bound NiRAN structure (Figure S8C) (Yan et al., 2021). In contrast to the AT-9010-bound NiRAN reported here, the diphosphate moiety of GDP was buried in the closed cavity formed by Lys50, Asn52, Lys73, and Arg 116, and was coordinated by a single Mg^2+^ ion (Figure 7). Similarly, the pseudo-kinase structures with a non-hydrolysable NTP show the γ-phosphate binding in a cavity formed by equivalent residues, with the ribose stabilized along a wide surface close to the groove (Figure S8C). It is thus reasonable to propose that AT-9010 has a unique binding mode, driven by both the hydrophobic nature of the cavity and the modified ribose. The base and ribose are stabilised in the cavity by conserved residues, accounting for a potent inhibition of the NiRAN function, consistent with enzymatic inhibition and thermal shift data.

**Figure 7.**
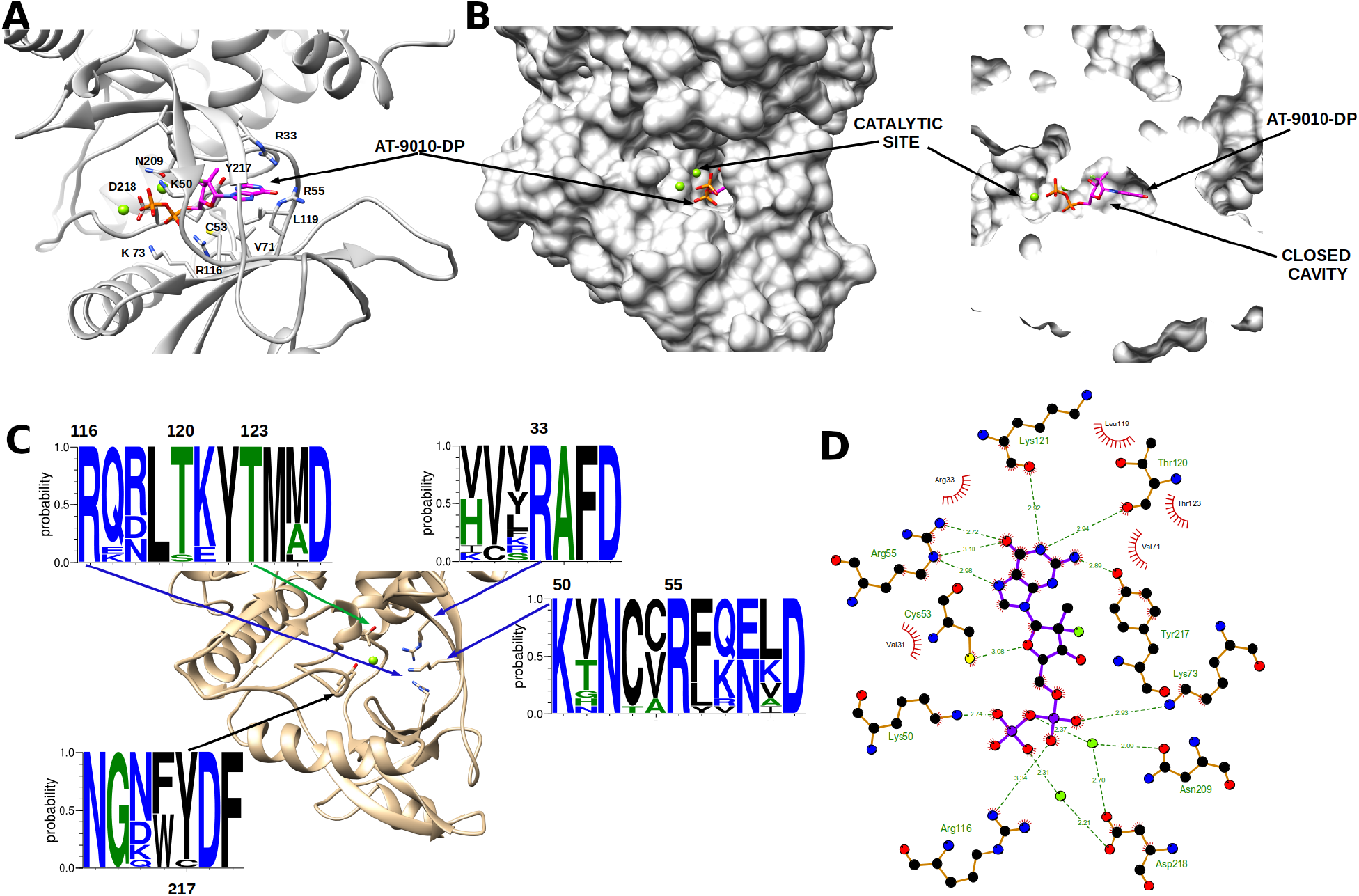
Binding of AT-9010 5’-diphosphate in the NiRAN domain. (A) Zoom of AT-9010 in its diphosphate form binding to NiRAN domain (Ribbon and sticks). (B) Left panel surface representation of the NiRAN showing AT-9010 engulfed in the cavity and 2 phosphate sticking out. Right panel sliced representation of the binding cavity with AT-9010 5’-diphosphate. (C) Coronavirus Weblogo of the NiRAN conserved residues around the cavity. (D) Ligplot 2D representation of the detailed interactions of AT-9010 binding in the NiRAN cavity.

## Discussion

Here, we show that the SARS-CoV RTC can initiate RNA synthesis through two independent pathways, mimicking viral (-)RNA synthesis. Both pathways are sensitive to AT-9010, and to a lesser extent STP, supporting a new antiviral mechanism for these nucleotide analogues.

It is intriguing that two separate mechanisms operate to launch synthesis of the (-)RNA strand. The preferential initiation at either a poly(A) sequence (protein-primed) or at the genome-poly(A) junction via pppGpU dinucleotide synthesis (*de novo*) suggests these two pathways are not redundant. Rather, this may serve as a mechanism to regulate continuous *vs* discontinuous viral RNA synthesis, for generation of full-length genomic RNA and sgRNAs, respectively. Of note, for (+)RNA viruses, cellular levels of positive-sense RNA are significantly higher than that of negative-sense RNA. This may explain why nsp8-linked viral genomes have been difficult to detect, in contrast to *Picornaviruses* which use a VPg to prime both plus and minus strand synthesis. Alternatively, we cannot exclude the possibility that nsp8-UMPylation and subsequent protein-priming is used in viral factories to initiate synthesis of poly(U)-containing sequences complementary to polyadenylated cellular mRNAs, thus serving as an immune decoy mechanism.

The UMPylation of nsp8 and resulting protein-primed pathway represents a first convincing role for the transferase-related activity of the NiRAN domain. Given its structural relatedness to the SelO family of pseudo-kinases, including YdiU which was recently shown to UMPylate up to 56 different peptides in response to stress (Sreelatha et al., 2018), it seems plausible that NiRAN-mediated UMPylation of other viral, and potentially host proteins may serve additional roles in the viral life-cycle. The labeling of nsp8 at different and/or multiple sites may additionally serve as a regulatory mechanism for its activity. Nsp8 is multimeric in solution, and additionally is known to form hexamers with nsp7. Differential UMPylation sites and levels may therefore serve to modulate its activity, as is the case for the *E.coli* glutamine synthetase enzyme, whose activity is dependent on the number of AMPylated subunits in the oligomeric structure (Itzen et al., 2011). The NiRAN domain has additionally been proposed to play a role in the capping pathway as a guanyltransferase (Yan et al., 2021), which may explain the increased stability provided by GTP binding compared with UTP, and the specificity for these two nucleotides. The preference for two distinct nucleobases (*i.e*., a purine and a pyrimidine) is not unheard of. For example, YdiU was recently shown to use both ATP and UTP as substrates, with self-AMPylation used as a mechanism to downregulate its UMPylation activity. In the case of CoVs, the switch between GTase and UMP-transferase activities may be regulated by other viral protein(s), such nsp9 (Slanina et al., 2021) which have also been reported to be UMPylated by nsp12. Of note, (Slanina et al., 2021) did not identify nsp8 as a target for NiRAN-mediated UMPylation, which we attribute to the presence of a C-terminal His-tag precluding nsp8-labeling.

Despite the preferential labeling of nsp8 by UTP, GTP is seen to be the more stably-bound substrate in the NiRAN active site. In accordance, inhibition assays and thermal shift data reflect the preferential binding of the guanosine analogue AT-9010 over its uridine equivalent STP. This is logical given the more transient UTP binding required to facilitate transfer to nsp8, and suggests that two NTP-binding modes exist, dependent on the nucleobase. It thus seems likely that guanosine NAs are the best candidates for NiRAN-based inhibition. Of note, ^m7^GTP did not inhibit nsp8-UMPylation, suggesting that more stringent selection occurs on the base rather than on the ribose. In the case of AT-9010, structural analysis shows that the 2’-fluoro-2’-C-methyl ribose modification is well accommodated in the NiRAN active site and provides additional interactions which likely contribute to its stability.

Furthermore, AT-9010 additionally targets the RdRp-domain for incorporation into viral RNA causing immediate chain termination, and impeding *de novo* synthesis initiation. Treatment of CoV-infections with NAs has been largely complicated by the presence of the ExoN activity. The pleiotropic action of AT-9010, in its dual targeting of NiRAN and RdRp domains, should attenuate the chance of resistance mutations. Our structural and functional description of its mechanism of action should aid further design of potent drugs, and be an important asset in the control of the expanding genetic pool of SARS-CoV2 variants observed in the current pandemic.

## Acknowledgements

This work was supported by the Fondation pour la Recherche Médicale (Aide aux Équipes), the SCORE project H2020 SC1-PHE-Coronavirus-2020 (grant#101003627), Inserm through the REACTing initiative (REsearch and ACTion targeting emerging infectious diseases), and ATEA, Inc. We thank Dr. Manfu Wang, and Dr. Zenglin Yuan from WuxiBiortus Biosciences Co. Ltd. for their help in data processing and refinement. We thank Minqi Gao from WuxiBiortus Biosciences Co. Ltd. for his help in enzyme activity assays, and Dr. Zheng Liu from the Centre for Electron Microscopy, CUHK, for his kind help in data collection. We are in debt to Pr. Olve Peersen for his constant support, data sharing, and valuable biochemical insight during the course of the project.

## Author contributions

V.F., B.Sa., M.F., C.F., P.G., P.E.K., C.E., M.N., performed experiments, E.D., K.A., N.R., J.-G., F.D., J.- J.V., S.G., K.L., J.-P.S. analyzed data, Y.Z., X.Y, H. S., performed experiments, collected and analyzed data, B.Se, A.M. performed experiments and analyzed data, F.F analyzed data and wrote the manuscript, A.S. and B.C. designed experiments, performed experiments, analyzed data, and wrote the manuscript.

## Declaration of interest

S.G., A.M., K.L. and J.P.S. are employees of ATEA Pharmaceuticals, Inc. The other authors declare no competing interests.

## Material and Methods

### 5’-triphosphate nucleosides

AT-9010, 5’-triphosphate 2’-fluoro-2’-C-methyl uridine, and Remdesivir 5’-triphosphate were from NuBlocks LLC, Oceanside, CA, USA. Other NTPs were purchased from GE Healthcare. 5’-TP dinucleotides were purified by semi-preparative IEX-HPLC with a UHPLC Thermoscientific Ultimate 3000 system equipped with a HPG-3200 BX pump, a DAD 3000 detector, a WPS-3000TBRS Autosampler, a fraction collector F, using a DNAPac PA200 column (22 × 250 mm). Elution was performed with buffer A: 5% CH_3_CN in 25 mM Tris-HCl pH 8 and buffer B: 5% CH_3_CN containing 400 mM LiClO_4_ in 25 mM Tris-HCl pH 8 at a 9 mL.min-1 flow rate. The crude dinucleotides were purified using a 0 - 15% linear gradient of buffer B in buffer A in 25 min at 25°C. The pure fractions were pooled in a 100 mL round bottomed flask and were evaporated under reduced pressure with a bath at 30°C maximum. The residue was desalted using a C_18_ cartridge Sep-Pak^®^ Classic. The residue was dissolved in 1.2 mL water (divided in 3 portions of 0.4 mL for flask rinse), transferred to a 2 mL Eppendorf-vial and lyophilized from water. Pure 5’-TP dinucleotides were characterized by MALDI-TOF mass spectrometry using a Axima Assurance spectrometer equipped with 337 nm nitrogen laser (Shimadzu Biotech, UK) using a 10:1 (m/m) mixture of 2,4,6- trihydroxyacetophenone/ammonium citrate as a saturated solution in acetonitrile/water (1:1, v/v) for the matrix. Analytical samples were mixed with the matrix in a 1:1 (v/v) ratio, crystallized on a 100-well stainless-steel plate and analyzed. UV quantification of 5’-TP dinucleotides was performed on a UV-1600 PC spectrometer (VWR) by measuring absorbance at 260 nm.

### Dinucleotide synthesis

Chemical synthesis of 5’-triphosphate (TP) dinucleotides pppUpU, pppUpG, pppGpU and pppApU were performed on an ABI 394 synthesizer (Applied Biosystems) from long chain alkylamine controlled-pore glass (LCAA-CPG) solid support with a pore size of 1000 Å derivatized through the succinyl linker with 5’-*O*-dimethoxytrityl-2’-*O*-acetyl-[uridine or *N2*-isopropylphenoxyacetyl guanosine] (Link Technologies). Dinucleotides were assembled on a 1-μmole scale in Twist oligonucleotide synthesis columns (Glen Research) using the 5’-*O*-DMTr-2’-*O*-pivaloyloxymethyl-3’-*O*-(*O*cyanoethyl-*N,N*-diisopropylphosphoramidite)-[uridine or *N*2-isopropylphenoxyacetyl guanosine or *N6*-phenoxyacetyl adenosine] (Chemgenes). After assembly, the CPG beads were dried under a stream of argon. A solution (2 mL) of 1 M diphenyl phosphite (0.4 mL) in dry pyridine (1.6 mL) was passed manually with a plastic syringe through the Twist column and left to stand for 30 min at 40°C. The CPG was then washed with acetonitrile and a 0.1 M solution of triethylammonium bicarbonate (TEAB, pH 7.5) was applied to the column and left to react for 45 min at 40°C. After several washings, an oxidation solution containing imidazole (150 mg) in *N,O*-bis-trimethylsilylacetamide (0.4 mL), acetonitrile (0.8 mL), bromotrichloromethane (0.8 mL) and triethylamine (0.1 mL) was added under argon and left to react for 2 h at 40°C. After washing and drying the support, a solution containing bis(tri-*n*-butylammonium) pyrophosphate (88 mg, 0.15 mmol) in dry DMF (0.5 mL) was applied to the column and left to react for 18 h at 40°C. The solution was removed, and the support was washed with dry acetonitrile. The CPG beads were dried by blowing argon through them for 1 min. 5’-TP dinucleotides were deprotected and released from the solid-support using a 1 M solution of 1,8-diazadicyclo-[5,4,0]undec-7-ene (DBU) (0.3 mL) in anhydrous acetonitrile (1.7 mL) for 3 min then the CPG beads were transferred into a glass vial and a 30 % aqueous ammonia solution (2 mL) was applied for 3 h at 40°C. The ammonia solution was collected in a 100 mL round bottomed flask and was evaporated and co-evaporated with water under reduced pressure with a bath at 30°C maximum. The residue was dissolved in water (1.8 mL divided into four portions for flask rinse: 0.6 mL, 0.4 mL, 0.4 mL, 0,4 mL), transferred to 2 mL Eppendorf-vials and then lyophilized from water.

### Expression and purification of SARS-CoV proteins

SARS-CoV cofactor proteins nsp7(TEV)6His, 6His(TEV)nsp8 and nsp7L8(TEV)6His were expressed under the control of a T5-promoter in pQE30 vectors in *Escherichia coli* (*E. coli*) NEB Express C2523 cells (New England Biolabs) carrying the pRARE2LacI (Novagen) plasmid. Protein was expressed overnight at 17°C (with 100 μg/mL Ampicillin and 17 μg/mL Chloramphenicol), following induction with 100 μM IPTG at an OD_600_ = 0.5 – 0.6. Cells were lysed by sonication in lysis buffer (50 mM Tris-HCl pH 8, 300 mM NaCl, 10 mM Imidazole, supplemented with 20 mM MgSO_4_, 0.25 mg/mL Lysozyme, 10 μgg/mL DNase and 1 mM PMSF) and protein was purified through affinity chromatography with TALON® Superflow™ cobalt-based IMAC resin (Cytiva). A wash step with buffer supplemented with 500 mM NaCl was performed prior to elution with 200 mM imidazole. The affinity tag was removed via overnight cleavage with TEV protease (1:10 w/w ratio to TEV:protein) in a dialysis buffer containing no imidazole and supplemented with 1 mM DTT. Cleaved protein was re-purified through a second cobalt column to remove the histidine-labelled TEV protease, and further purified with size exclusion chromatography (Cytiva Superdex S200) in a final buffer of 25 mM HEPES pH 8, 150 mM NaCl, 5 mM MgCl_2_ and 5 mM TCEP. SARS-CoV nsp12-8His was expressed from a pJ404 vector in *E. coli* strain BL21/pG-Tf2 (Takara 9124), in the presence of Ampicillin (100 μM/mL) and Chloramphenicol (17 μg/mL). Expression was induced at OD_600_= 0.5-0.6 with 250 μM IPTG and 5 ng/ml of tetracycline for induction of chaperones proteins (groES-groEL-tig), and left overnight at 23°C at 220 rpm. Cells were lysed over 45-60 mins at 4°C, in a buffer containing 50 mM Tris pH 8, 300 mM NaCl, 5 mM MgSO4, 10% glycerol, 1% CHAPS, supplemented with 5 mM 2-mercaptoethanol, 0.5 mg/ mL Lysozyme, 10 μgg/mL Dnase, 1 mM PMSF, 0.2 mM Benzamidine. Nucleic acid was precipitated through the gradual addition of NaCl to a final concentration of 1 M. Following centrifugation (30000 × g for 30min), the supernatant was diluted to reduce NaCl concentration to 300 mM. Protein was purified using cobalt-based IMAC resin TALON® Superflow ^™^ (Cytiva), washing 3 times with wash buffer (50 mM Tris pH 8, 10% glycerol) with alternating NaCl concentration (300 mM, 1 M, 300 mM) before elution with 200 mM Imidazole. Protein was further purified through size exclusion chromatography (Cytiva Superdex S200) in a the same final buffer as 7 and 8 cofactor proteins, supplemented with 10% glycerol. Concentrated aliquots of nsp12, 7, 8 and 7L8 were flash-frozen in liquid nitrogen and stored at −80 ° C. SARS-CoV cofactor protein nsp10 was expressed under the control of a Tet-promoter in a pASK vector in *E. coli* NEB Express C2523 cells (New England Biolabs) carrying the pRare2LacI (Novagen) plasmid. Protein was expressed overnight at 17°C (with 50 μM/mL kanamycin, 17 μg/mL Chloramphenicol), following induction with 200 μg/L Tetracycline at an OD_600_ = 0.6-0.7. Cells were incubated in lysis buffer (50 mM HEPES pH 7.5, 300 mM NaCl, 10 mM Imidazole, 5 mM MgSO_4_, 1 mM of β-mercaptoethanol, 0.25 mg/mL Lysozyme, 10 μgg/mL DNase, 0.1% triton and 1 mM PMSF) and lysed by sonication. Protein was purified first through affinity chromatography with HisPur Cobalt resin (Thermo Scientific), eluted in 100 mM imidazole, then through size exclusion chomatography (GE Superdex S200) in a final buffer of 50 mM HEPES pH 7.5, 300 mM NaCl, 5 mM MgCl_2_ and 1 mM of β-mercaptoethanol. SARS-CoV nsp14 was expressed from a pDEST14 vector in *E. coli* strain NEB Express C2566 cells (New England Biolabs) carrying the pRare2, in the presence of Ampicillin (100 μM/mL) and Chloramphenicol (17 μg/mL).Protein expression was induced at an OD_600_ = 0.8 with 2 μM IPTG, and left overnight at 17°C with shaking. Cells were lysed by sonication in a buffer containing 50 mM HEPES pH 7.5, 500 mM NaCl, 20 mM Imidazole, supplemented with 0.25 mg/mL Lysozyme, 10 μgg/mL DNase and 1 mM PMSF. The protein was purified through affinity chromatography with HisPur Cobalt resin (Thermo Scientific), washing with an increased concentration of salt (1 M NaCl), prior to elution in buffer supplemented with 250 mM imidazole. The protein was further purified by a size exclusion chomatography (GE Superdex S200) in a final buffer of 10 mM HEPES pH 7.5, 150 mM NaCl.

### Expression and purification of SARS-CoV-2 proteins

SARS-CoV2 proteins were either purchased from Biortus (en.wuxibiortus.com) or purified for Cryo-EM using the following protocol. The gene of the full-length SARS-CoV-2 nsp12(residues 1-932) was synthesized with codon optimization (General Biosystems). and cloned into pFastBac1 baculovirus expression vector. An additional peptide (HHHHHHHHWSHPQFEKENLYFQG) was added to the N-terminus of nsp12. *Spodoptera frugiperda* (Sf21) cells expressing the target protein were collected 48 h after infection at 27 °C and were centrifugation at 4,500 rpm for 10 min. Pellets were resuspended in lysis buffer (50mM Tris-HCl (pH 8.0), 500mM NaCl, 5% glycerol, 2mM MgCl_2_, cOmplete Protease Inhibitor Tablet) and homogenized with High Pressure Homogenizer at 4℃. Cell lysate was centrifuged at 18,000 rpm for 60 min at 4℃. The fusion protein was first purified by Strep-Tactin (Strep-Tactin®XT) affinity chromatography and the tag was removed by incubation of TEV protease overnight at 4°C after elution. The protein was reloaded onto a Heparin HP column after buffer exchanged to buffer A (50mM Tris-HCl, pH8.0, 150mM NaCl, 5% glycerol, 2mM MgCl_2_). Flow through was collected and loaded on to a HiLoad 16/600 Superdex 200 pg column (GE healthcare) equilibrated in 10 mM Tris-HCl, pH 8.0, 500 mM NaCl, 2 mM MgCl_2_. Purified nsp12 was concentrated to 6.86 mg/ml and stored at −80°C. The gene of SARS-CoV-2 nsp7 (residues 1-83) possessing a C-terminal Avi-6Histag (GLNDIFEAQKIEWHEHHHHHH) was cloned into a modified pET-32a vector. BL21(*E. coli*, T7 Express) containing the plasmid were grown to an OD_600_ of 0.6 at 37 °C, and protein was expressed at 15 °C for 16 h after the addition of isopropyl β-D-1-thiogalactopyranoside (IPTG) to a final concentration of 0.5 mM. The cells were harvested then resuspended in buffer B (50mM Tris-HCl (pH 8.0), 500mM NaCl, 5% glycerol, 10mM imidazole). Cells were disrupted by a High-Pressure Homogenizer at 4°C. The insoluble material was removed by centrifugation at 18,000 rpm, 60 min at 4°C. The fusion protein was purified by Ni-NTA (Novagen, USA) affinity chromatography followed by a Superdex HiLoad 16/600 Superdex 75 pg column (GE Healthcare, USA) in buffer C (10mM Tris-HCl (pH 8.0), 150mM NaCl). Purified nsp7 was concentrated to 7.27mg/ml and stored at −80°C. The gene of SARS-CoV-2 nsp8 (residues 1-198) was cloned into a modified pET-28a vector containing an N-terminal His6-flag-tag with a TEV cleavage site (HHHHHHDYK DDDDKENLYFQG) for expression in *E. coli*. Nsp8 was expressed in the same way as that for nsp7. Cells were harvested then resuspended in buffer D (50mM Tris-HCl, pH 8.0, 500mM NaCl, 5% glycerol). Cells were lysed using a High-Pressure Homogenizer at 4 °C. Cell lysate was clarified by centrifugation at 18,000 rpm, 60 min at 4°C. Supernatant was applied onto a Talon affinity chromatography column and tag was removed by on-column cleavage overnight using TEV protease. The mixture was buffer exchanged to buffer D and reloaded onto a His-chelating NTA column again to remove the His tag and TEV protease. Target protein was further purified by passage through a HiLoad 16/600 Superdex 75 pg (GE Healthcare, USA) in buffer E (20 mM Tris-HCl, pH 8.0, 200 mM NaCl, 5% glycerol). The fractions near the maximum height of the peak were combined and further purified by a Mono Q 10/100 GL column (GE Healthcare, USA). Buffer exchange to buffer E. Purified nsp8 was buffer exchanged to buffer E, then concentrated to 11.63 mg/ml and stored at −80 °C.

### Thermal shift assays

The influence of compound binding on protein stability was measured by Themofluor assay, using a CFX Connect BioRad real-time PCR machine, at a protein concentration of 2 μM (nsp12 WT and mutants) in a buffer composed of 10 mM HEPES pH7.4, 150 mM NaCl and 5 mM MgCl_2_ supplemented or not by 0.5 mM MnCl_2_ freshly prepared. Compounds and NTPs concentration was fixed at 100 μM. In 96-well thin-walled PCR plates, 2 μL of inhibitors or NTPS were added to 16 μL of buffer follow by the addition of 2 μL of protein. Finally, 2 μL of the fluorescent dye SYPRO Orange was added (5-fold final). Melting-temperature (Tm) values given are the average and standard deviation of three independent experiments.

### Nucleotidyl transferase reaction

Standard nucleotide transfer reactions were performed at 37°C for reaction times ranging from 2 – 60 min, in a buffer containing 20 mM HEPES pH 7.5, 1 mM DTT, 0.5 - 2 mM MnCl_2_, 1 – 5 μCi α^32^P-NTP and 30 mM NaCl with final protein concentrations of 1 μM nsp12 and/or 3 μM cofactors. For analysis of labeling only, samples were stopped in a 2X concentration of SDS-loading dye and heated at 95°C for 5 mins, to ensure only covalently-bound NMP remained bound to protein. Proteins were analyzed on 15% SDS PAGE gels, stained with InstantBlue for total protein and exposed for 2 hrs – overnight to reveal radiolabeled proteins. For assessment of nsp8-UMP bond stability, nsp8 was first labeled in a reaction containing only nsp12 protein, or with the full RTC comprised of nsp12:nsp7L8:nsp8 in the presence of RNA. Reactions were heat inactivated at 70°C for 10 mins, then incubated with either 0.1M HCl or 0.1M NaOH for 1 hour at 30°C. Following incubation, the pH was neutralized with an equivalent concentration of acid or base, and samples were analyzed by SDS PAGE, as above. Competition experiments with AT-9010 and STP were run in the same conditions as above, at two enzyme concentrations, 1 μM or 5 μM of nsp12 with 5-fold molar excess of nsp8. A constant concentration of either GTP or UTP (5 μM), supplemented with 5 μCi α^32^P-NTP was used, with varied concentrations of NA inhibitor. Reactions were stopped after either 15 min – 1 hr incubation, and analyzed on 15% SDS PAGE gels. Nsp8-UMPylation or GMPylation was calculated as a percentage relative to the no-inhibitor controls. Reactions were performed in triplicate with 6-7 concentrations of NA.

### Primer-independent polymerization assays

Poly(N)27 and oligonucleotide substrates corresponding to the 3’ end of the SARS-CoV genome (with or without a poly(A)_15_ tail) were purchased from Biomers (HPLC grade). The active RTC was formed by first incubating nsp7 and 8 together at equimolar concentrations (100 μM) for 30 mins at room temperature. Nsp12, extra nsp8 and protein gel filtration buffer were added to form a final complex consisting of nsp12:7:8 at a 1:3:6 ratio, with 10 μM nsp12. The complex was further incubated for 10 mins at room temperature, and used at a final concentration of 1 μM nsp12. Primer-independent assays were performed in the same conditions used for nucleotidyl transferase reactions, but supplemented with 100 – 200 μM cold NTP and 0.7 μM final concentration of RNA. Reactions were stopped at indicated time-points in either 2X concentration of SDS loading buffer for protein analysis, or a 2X-4X volume of FBD stop solution (formamide, 10 mM EDTA) for analysis on 14% acrylamide, 7M urea sequencing gels. For proteinase K digestions, reactions were first stopped by heat-inactivation at 10 mins at 70°C. Digestion was performed for 2 hours at 37°C in a buffer containing 20 mM HEPES pH 7.5 and 0.5 % SDS. Nuclease P1 (NEB) were performed as per manufacturer’s instructions. For order of addition experiments, protein was pre-incubated in reaction buffer for 30 min at 37 °C with either UTP or RNA. Following incubation, the inverse reagent was added to start the reaction. AT-9010 and STP inhibition was tested with a constant concentration of 200 μM cold NTP and varied concentrations of inhibitor. Inhibitor, NTP and RNA were incubated together in assay buffer, and reactions were started via addition of the protein complex. For analysis of covalent binding to nsp8 with western blotting, polymerase assays were performed in the buffer described above. Nsp12:7:8 complex was preincubated with 200 μM UTP for 30 min at 37°C prior to addition of 2 μM poly(A)_27_, ST20poly(A)_15_ or ST20 RNA. Reactions were either stopped immediately after RNA addition (time 0), or after 60 min incubation at 37°C in 2X SDS loading buffer. Samples were separated on 15% SDS-PAGE gels, transferred to nitrocellulose membranes and blocked in TBS-Tween (0.1%) containing 5% skim milk powder overnight. Nsp8 was probed with mouse monoclonal anti-nsp8 (5A10) from GeneTex (GTX632696), and HRP-conjugated rabbit anti-mouse secondary (Agilent Dako), washing 3X in PBS.T between each antibody. Nsp8 was revealed with Immobilon Crescendo Western HRP Substrate (Millipore).

### Primer-dependent polymerization and excision assays

Primer-template (10:20) pairs, corresponding to the 3’ end of the SARS-CoV genome, were annealed at a molar ratio of 1:1.5 in 110 mM KCl at 70 °C for 10 min, then cooled slowly to room temperature over several hours. Hairpin RNAs were synthesized by Integrated DNA Technologies (Coralville, IA). The nsp12:7:8 RTC was formed as for the primer-independent polymerase assays, then preincubated with RNA in a pre-mix containing 20 mM HEPES pH 7.5, 50 mM NaCl, 5 mM MgCl_2_. For single nucleotide incorporation assays, reactions were initiated with 50 μM (final concentration) of all AT-9010 or STP, with or without the following nucleotide (ATP). Final reaction concentrations were 0.5 μM nsp12, 0.4 μM RNA. Reactions were quenched at indicated timepoints with 5X volume of FBD stop solution (formamide, 10 mM EDTA). For AT-9010 – GTP competition experiments, protein-RNA complexes were preincubated as described above, and initiated with either all four NTPs, or with only CTP, UTP and ATP (50 μM each NTP), supplemented with various concentrations of AT-9010 (10 – 250 μM). To calculate the discrimination between AT-9010 and GTP, the AT-9010 product band (from 50 or 250 μM concentrations) was compared with the sum of fractions of product bands derived from GTP incorporation at three timepoints. Discrimination was corrected to account for concentration difference between AT-9010 and GTP. For analogue excision, polymerization reactions were performed on hairpin RNAs in the same conditions as described above, then stopped by heating at 70°C for 10 min. The hairpin was re-annealed at 30° for >30 min, and 50 nM nsp14/nsp10 (1:5) were added for time-course reactions. Aliquots were analyzed using denaturing polyacrylamide gel electrophoresis (20 % acrylamide, 7M urea) and visualized using a Typhoon FluorImager.

### Assembly of the extended nsp12-nsp7-nsp8-RNA complex for Cryo-EM

To assemble the extended RdRp complex, nsp12 was incubated with nsp7 and nsp8 at 4 °C for 3 h with a molar ratio of 1: 3: 6 in a buffer containing 25 mM Tris-HCl (pH 8.0), 50 mM NaCl, 5 mM MgCl_2_, 4 mM DTT. Then the mixture was purified by Mono Q 5/50 GL ion-exchange chromatography (GE Healthcare, USA), resulting in a stable nsp7-nsp8-nsp12 complex. The protein complex was desalted to buffer F (25mM Tris-HCl (pH 8.0), 100 mM NaCl, 5 mM MgCl_2_, 4 mM DTT). Purified RdRp complex were buffer exchanged to 25 mM Tris-HCl, pH 8.0, 100 mM NaCl, 5 mM MgCl_2_, 4m M DTT and concentrated to 10 mg/ml for Cryo-EM experiments. A 30-mer oligoribonucleotide template and a 20-mer oligoribonucleotide primer were chemically synthesized by GenScript. The template and primer oligoribonucleotides were annealed by heating the solution to 95 °C and gradually cooling to 4 °C. The annealed RNA scaffold was incubated with nsp12-nsp7-nsp8 complex for 30 min at 4 °C with a molar ratio of 2: 1 to form the nsp12-nsp7-nsp8-RNA complex. AT9010 was added subsequently for compound incorporation.

### Cryo-EM sample preparation and data collection

In total, 3 μL of protein solution at 5 mg/mL (with 0.025% DDM) was applied onto a glow-discharged holey carbon grid (Quantifoil, 300 mesh, R1.2/1.3). Excess samples were blotted for 5.0 s with a blotting force of 3, then the remaining solution was vitrified by plunging into liquid ethane using a Vitrobot Mark IV (Thermo Fischer Scientific) at 4 °C and 100% humidity. Cryo-EM data were collected with a 300 keV Titan Krios electron microscope (Thermo Fisher Scientific, USA) equipped with a K3 direct electron detector (Gatan, USA) operating in a super-resolution counting mode. All movies were automatically recorded using SerialEM (Mastronarde, 2005) at a magnification of 105K, with a physical pixel size of 0.83 Å. A total dose of 80.5 e^-^/ Å^2^ was fractionated into 50 frames. 7,459 movie micrographs were collected with a defocus range from −1.5 μm to −2.5 μm, and the slit width of Gatan Quantum GIF energy filter (Gatan, USA) was set to be 20 eV. Statistics for data collection and refinement are shown in Table S1.

### Cryo-EM image processing

All dose-fractioned movies were motion-corrected with Relion’s own implementation. CTF estimation, 2D classification, 3D classification and refinements were all performed in cryoSPARC. A total of 2,410,466 particles were auto-picked using blob picker and extracted with a box size of 320 pixels. 248,401 particles were selected after three rounds of 2D classification based on the complex integrity. This particle set was used for Ab-Initio reconstruction with three classes, which were then used as 3D volume templates for heterogeneous refinement. The 3D volume corresponding to the intact nsp12-nsp7-nsp8-RNA complex was used for creating 100 2D projections which were then used as templates for template-based particle picking. Approximately 3,640,595 particles were picked from a set of 3,622 micrographs filtered based on fitted resolution better than 5 Å as estimated by CTFFIND4, using template picker. Particles were extracted with a box size of 360 pixels. A total of 234,421 particles were selected after four rounds of 2D classification based on the complex integrity. This particle set was used for Ab-Initio reconstruction with three classes, followed by heterogeneous refinement. A subset of 181,669 particles from the class with good features was subjected to Homogeneous Refinement, Local Refinement and Non-uniform Refinement, resulting in a 2.98 Å map.

### Model building and refinement

To build the model of nsp12-nsp7-nsp8-RNA complex, the structure of SARS-CoV-2 nsp12-nsp7-nsp8-RNA complex (from PDB 7CYQ with one nsp9 and nsp13 removed) was placed and rigid-body fitted into the Cryo-EM map using UCSF Chimera. The model was manually built in Coot (Emsley et al., 2010) with the guidance of the Cryo-EM map, and in combination with real space refinement using Phenix (Afonine et al., 2018). The model validation statistics are shown in Table S1.

## Supplemental information

**Figure S1. Labeling of nsp8 by nsp12 WT and NiRAN mutants, in the presence of MnCl_2_ and MgCl_2_.**(A-B) SARS-CoV nsp12 NiRAN-mediated labeling of nsp8 with α^32^P-UTP in the presence of varied concentrations of MnCl_2_, MgCl_2_ or both ions. Samples were analyzed on 15% SDS PAGE and stained for total protein (top) and exposed to reveal covalently-bound radioactive UMP (bottom gels). A) Reactions performed with nsp12 and nsp8 in the absence of RNA, b) Reactions performed with nsp12:7L8:8 RTC in the presence of poly(A)_27_ RNA. (C) As in (B) but analyzed on 14% acrylamide urea PAGE gels. (D,E) Labeling of nsp8 with α^32^P-UTP (D) and α^32^P-GTP (E) was performed with the nsp12:7:8 complex with various NiRAN alanine mutants. Labeling reactions were performed for 1h at 37°C. (F) Denaturing SDS-PAGE analysis of SARS-CoV-2 RTC with α^32^P-UTP. Lane 1; without RNA, lane 2; protein complex pre-incubated with UTP + α^32^P-UTP prior to addition of poly(A)_27_ RNA, lane 3; protein complex pre-incubated with poly(A)_27_ RNA prior to addition of UTP + α^32^P-UTP, lanes 4-5; as in lane 2 and 3 but digested with PK

**Figure S2. Poly(U)_n_ synthesis from poly(A)_27_ RNA template by the RTC.** (A) Various combinations of SARS-CoV nsp12, nsp7 and nsp8, as well as a covalently linked version of nsp7 and 8 (nsp7L8) were incubated for 1hr at 37°C with UTP (supplemented with α^32^P-UTP) and poly(A)_27_ RNA. Samples were analyzed on a 14% acrylamide 7M urea PAGE sequencing gel, and correspond to samples shown in Figure 2a. (B) Activity of the nsp12:7:8 complex with three RNA substrates; i) poly(A)_27_ template RNA, ii) poly(A)_27_ template RNA blocked at the 3’ end via replacement of the 3’OH with a phosphate group (poly(A)_27_-3’P), and iii) poly(A)_27_ template labeled at the 5’ end with ^32^P (^32^P-poly(A)_27_). Synthesis was measured via addition of UTP supplemented with α^32^P-UTP for the first two RNAs, and with cold UTP only for ^32^P-poly(A)_27_. C represents reaction control, while PK shows same sample following proteinase K digestion. (C) Denaturing SDS-PAGE western blot analysis with anti-nsp8 (5A10). The RTC was preincubated for 30 mins with UTP prior to addition of different RNAs and remaining NTPs (if indicated); poly(A)_27_ (lanes 1, 2, 11), ST20poly(A)_15_ (lanes 3-6, 12, 13), ST20poly(A)_15_ blocked at the 3’ terminus with cy3 (lanes7-8), ST20 (lanes 9, 10, 14). Reactions were stopped immediately after RNA addition (time 0) or after 60 min incubation. Red line shows presence of nsp8-p(U)_n_ products with both poly(A)_27_ and ST20poly(A)_15_ when only UTP is given (lanes 2 and 6, respectively). Blue arrows show nsp8 covalently bound to full-length ST20poly(A)_15_ products. d-e) Activity of the RTC with poly(A)_27_, poly(U)_27_ or poly(C)_27_ RNA templates, with UTP, ATP or GTP (supplemented with the corresponding α^32^P-NTP), respectively. (D) Analysis of products using 7M urea PAGE. From the bottom up, solid dots show tri-, di- and mono- phosphates of uridine (red), adenosine (yellow) and guanosine (green), respectively. Asterixes show pppNpN dinucleotide products with same colour scheme. Synthesis of poly(A)_n_ from poly(U)_27_ template is characteristic of terminal transferase activity, potentially attributable to nsp8 (Tvarogová et al., 2019). (E) Analysis of products on 15% SDS PAGE gels before (left panel) and after treatment with proteinase K (middle panel), or nuclease P1 (right panel). Top panels show gels following exposure for radioactivity, bottom panels are same gels stained for total protein. Black arrow shows nsp8 position.

**Figure S3. Order of addition experiments with various NiRAN mutants.** (A) Activity of the RTC on poly(A)_27_ template RNA with either wild-type (WT) nsp12, or various NiRAN active site mutants. Red arrow shows protein-primed product. (B) Denaturing SDS-PAGE analysis of order of addition experiment corresponding to Figure 2C. The RTC was incubated for 30 min at 37 °C with either UTP or RNA. Following incubation, the complementary reagent was added and reactions were stopped at indicated timepoints. (C) Timecourse showing activity of various NiRAN mutant RTCs, in addition to SAA RdRp mutant. Complexes were preincubated with UTP prior to RNA addition. PK represents 60 min timepoint, digested with proteinase K. (D) Conditions as in b, comparing the activity of WT and mutant RTCs through order of addition experiments analyzed through denaturing SDS-PAGE. (E) Quantification of protein-primed activity over time (left) and residual protein-primed activity of NiRAN mutants compared to WT (right), based on red arrow, panel C. (F) Quantification of *de novo* synthesis activity over time (left), and fold-increase in p(U)_54_ product relative to WT (right), based on p(U)_54_ product in panel C.

**Figure S4. The nsp12:7:8 complex additionally mediates synthesis of pppNpN to prime synthesis.** (A) Synthesis of poly(U)_n_ RNA by the RTC from a poly(A)_27_ template was assessed in the presence of varying concentrations of pppUpU (0 – 100 μM) over time. (B) Quantification of p(U)_54_ (top) and pppUpU (bottom) products was performed with ImageQuant analysis software and plotted as a function of time. (C) Synthesis of heteropolymeric RNA from templates with and without a poly(A) tail. An RNA template mimicking the SARS-CoV-1 3’-end of the genome (ST20) extended by a poly(A)_15_ sequence (ST20pA_15_) was incubated with the RTC and NTPs supplemented with α^32^P-UTP. The same templates were 3’-blocked with a phosphate group (ST20-3’P) or a cy5 fluorescent dye (STP20p(A)_15_cy5), respectively. A control using a p(A)27 template is shown on right of gel. Size markers are shown as ST20 and ST20p(A)_15_ (rightmost lane). UMP, pppUpU, and UTP separated at the bottom of the gel are shown in correspondence to the top part of the gel. Synthesis products were separated using 7M urea PAGE (20 % acrylamide) and visualized using a Typhoon FluorImager.

**Figure S5. Incorporation and excision of AT-9010 and STP by the RTC.** (A) Guanosine analogue prodrug AT-511, and active metabolite AT-9010, in comparison to STP. (B) Incorporation of AT-9010 and STP as the first nucleotide with and without the following NTP. (C) Incorporation of AT-9010 in the presence (left) or absence of GTP (50 μM). Fold-preference for GTP > AT-9010 incorporation calculated by comparing the amount of AT-9010 insertion (red circle) relative to full-length product at two concentrations, at 3-timepoints. (D) Incorporation of AT-9010 and STP by the RTC (red circle), followed by excision with the nsp14 exonuclease (Exo).

**Figure S6. Labeling of nsp8 by nsp12 in presence of AT-9010, STP and m7GTP.** (A) Labeling of 5-fold molar excess of nsp8 by nsp12 (5 μM) was performed with a constant concentration of UTP (top) or GTP (bottom) supplemented with α^32^P-NTP (5 μM total), in competition with increasing concentrations of inhibitor or ^m7^GTP. Total intensity of labeling was quantified with ImageQuant software, and plotted at % residual activity. Calculated IC50 values were (B) 1.9 ± 0.1 and 8.2 ± 1.4 for AT-9010 and STP, respectively, in competition with UTP. (C) 2.7 ± 0.3 and 5 ± 0.8 for AT-9010 and STP, respectively, in competition with GTP. Data was calculated from two individual replic(Hartenian et al., 2020)ates, with a minimum of 6-points per replicate. (D) Inhibition of WT or K73A NiRAN mutant RTCs with varied concentrations of AT-9010 or STP, using both poly(A)_27_ and poly(C)_27_ templates in the presence of UTP and GTP, respectively (200 μM, supplemented with α^32^P-NTP).

**Figure S7. Cryo-EM data processing summary.** (A) The flowchart shows the main steps in data processing, from particle picking, through classification, to final maps. A selected subset of the initial reference-free 2D class averages and all the intermediate 3D class averages computed during the processing of this dataset are shown. All 3D class averages, selected for subsequent rounds of processing, are boxed in red, and the percentage of number of particles in each of these is shown. Further attempts to process the discarded classes are omitted from this chart for clarity, as these data did not contribute to the final particle set. (B-E) Map and model quality of the nps12-nsp7-nsp8/RNA/AT9010 complex. (B) FSC curves for the final reconstruction. The resolution of the map is 2.98 Å as calculated by the gold-standard FSC 0.143 criterion. (C) Viewing direction distribution of the final reconstruction. (D) The B-factor used for map sharpening is −82.7 Å^2^ as calculated by the Guinier Plot. (E) Model vs. map FSC curves. The resolution of the model is calculated to be 3.14 Å based on the masked map. (F) Local resolution distribution of the nps12-nsp7-nsp8/RNA/AT9010 complex calculated in cryoSPARC. Three views of the maps are shown rotated by 90° increments.

**Figure S8. Pseudo-kinase fold and conservation of the NiRAN domain of nsp12.** A) Sequence alignment derived from structural superimposition of several pseudo-kinase structure and NiRAN. (B) Sequence conservation (in blue) plotted onto the NiRAN domain structure. Conserved residues are in deep blue. (C) Electrostatic representations of NiRAN ions are in green spheres. Catalytic site goes from the ions toward the flat opening. Below the ions is the entry of the cavity. (D) Comparative analysis of nucleotide ligands into the NiRAN catalytic site with GDP (PDB: 7CYQ, left), AT-9010 (center), and ATP from the SelO structure (PDB: 6EAC, right), showing the different binding modes.

