## supplementary Figs for "Protein-primed RNA synthesis in SARS-CoVs and structural basis for inhibition by AT-527"

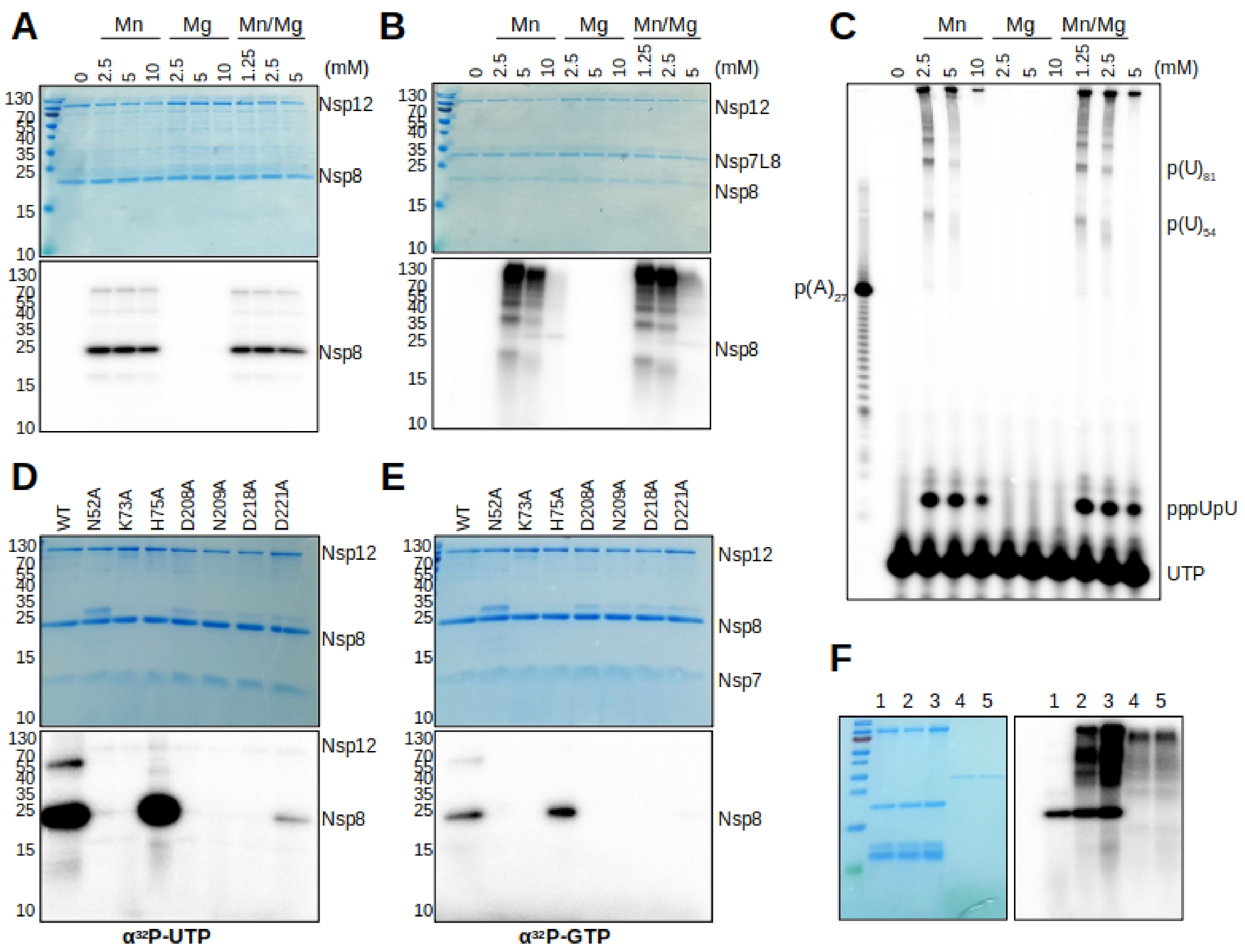
A
Mn
Mg
Mn/Mg
(mM)
Nsp12
Nsp8
B
Nsp7L8
C
p(U)81
p(U)54
p(A)27
pppUpU
UTP
D
WT
N52A
K73A
H75A
D208A
N209A
D218A
D221A
α32P-UTP
E
Nsp7
α32P-GTP
F

**Figure S1.**

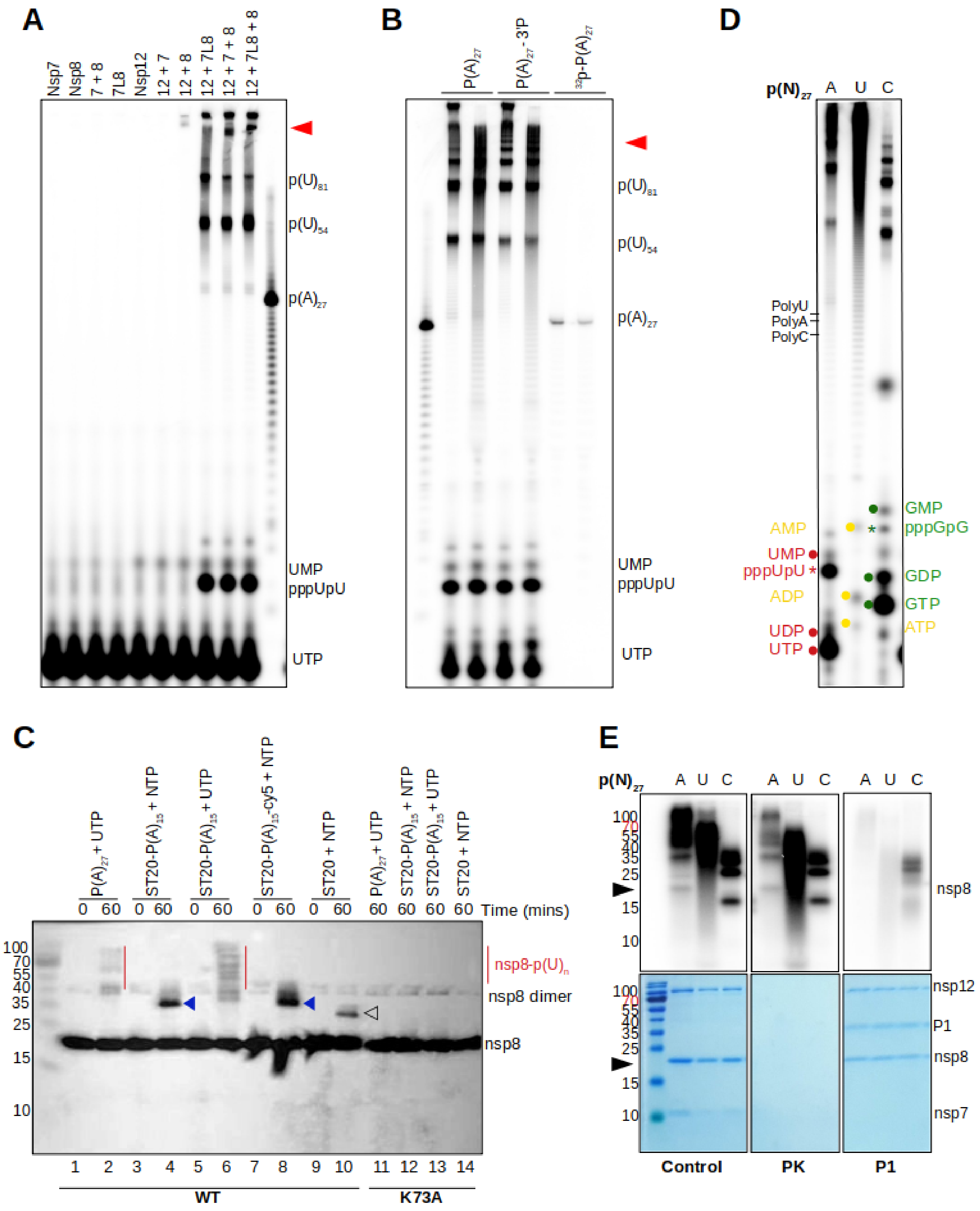
A
Nsp7
Nsp8
7 + 8
7L8
Nsp12
12 + 7
12 + 8
12 + 7L8
12 + 7 + 8
12 + 7L8 + 8
p(U)81
p(U)54
p(A)27
UMP
pppUpU
UTP
B
P(A)27
P(A)27 - 3'P
32p-P(A)27
D
p(N)27
A
U
C
PolyU
PolyA
PolyC
AMP
UMP
pppUpU
ADP
UDP
UTP
GMP
pppGpG
GDP
GTP
ATP
C
P(A)27 + UTP
ST20-P(A)15 + NTP
ST20-P(A)15 + UTP
ST20-P(A)15-cy5 + NTP
ST20 + NTP
Time (mins)
nsp8-p(U)n
nsp8 dimer
nsp8
WT
K73A
E
Control
PK
P1
nsp12
P1
nsp7

Figure S2.

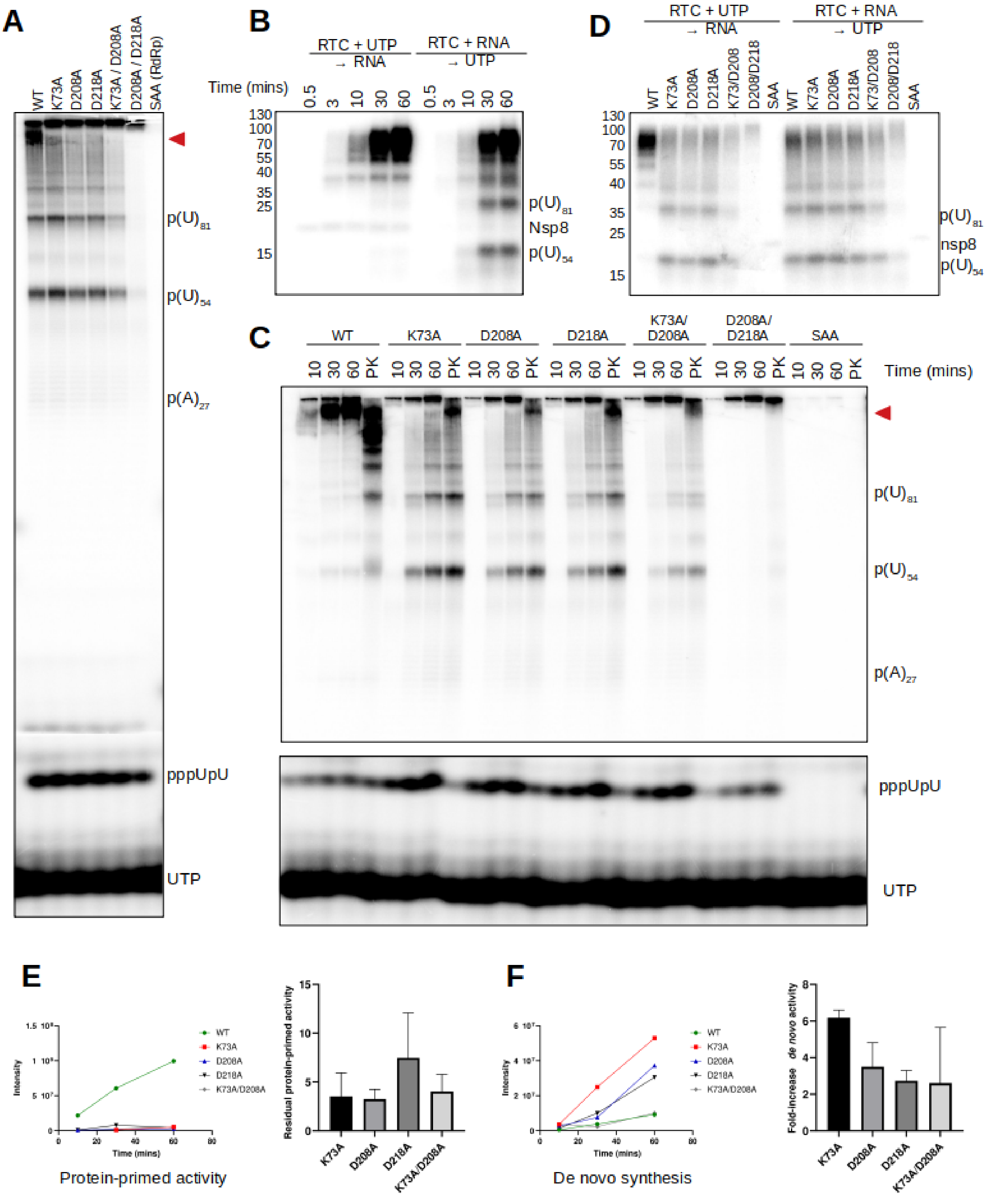
A
WT
K73A
D208A
D218A
K73A / D208A
D208A / D218A
SAA (RdRp)
p(U)81
p(U)54
p(A)27
pppUpU
UTP
B
RTC + UTP → RNA
RTC + RNA → UTP
Time (mins)
0.5
3
10
30
60
130
100
70
55
40
35
25
15
Nsp8
C
WT
K73A
D208A
D218A
K73A/ D208A
D208A/ D218A
SAA
10
30
60
PK
Time (mins)
D
K73/D208
D208/D218
nsp8
E
Protein-primed activity
Intensity
Time (mins)
Residual protein-primed activity
K73A/D208A
F
De novo synthesis
Fold-increase de novo activity

**Figure S3.**

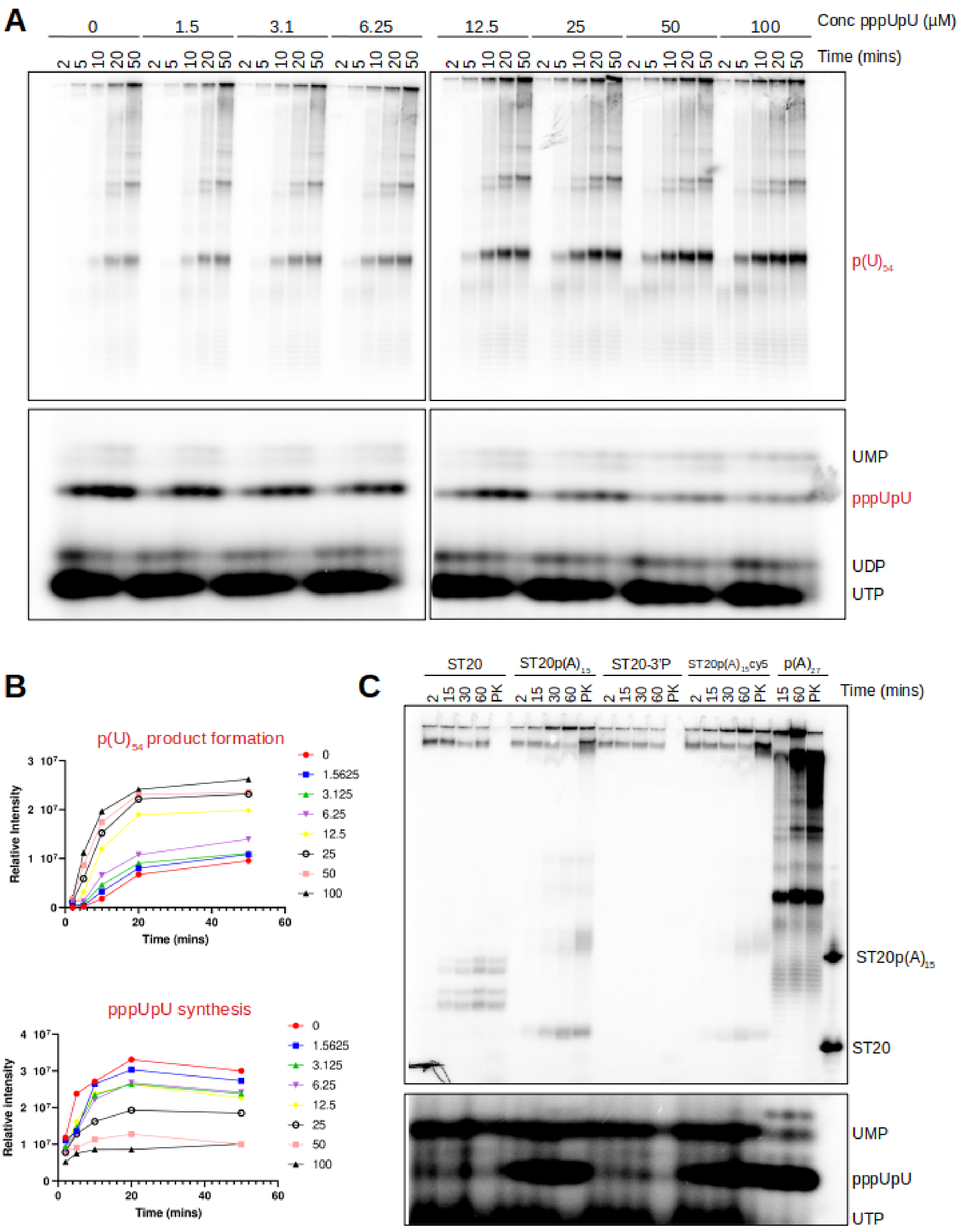
A
0
1.5
3.1
6.25
12.5
25
50
100
Conc pppUpU (µM)
2 5 10 20 50
Time (mins)
p(U)54
UMP
pppUpU
UDP
UTP
B
p(U)54 product formation
Relative Intensity
Time (mins)
0
1.5625
3.125
6.25
12.5
25
50
100
pppUpU synthesis
C
ST20
ST20p(A)15
ST20-3'P
ST20p(A)15cy5
p(A)27
2 15 30 60 PK
Time (mins)
ST20p(A)15
ST20
UMP
pppUpU
UTP

**Figure S4.**

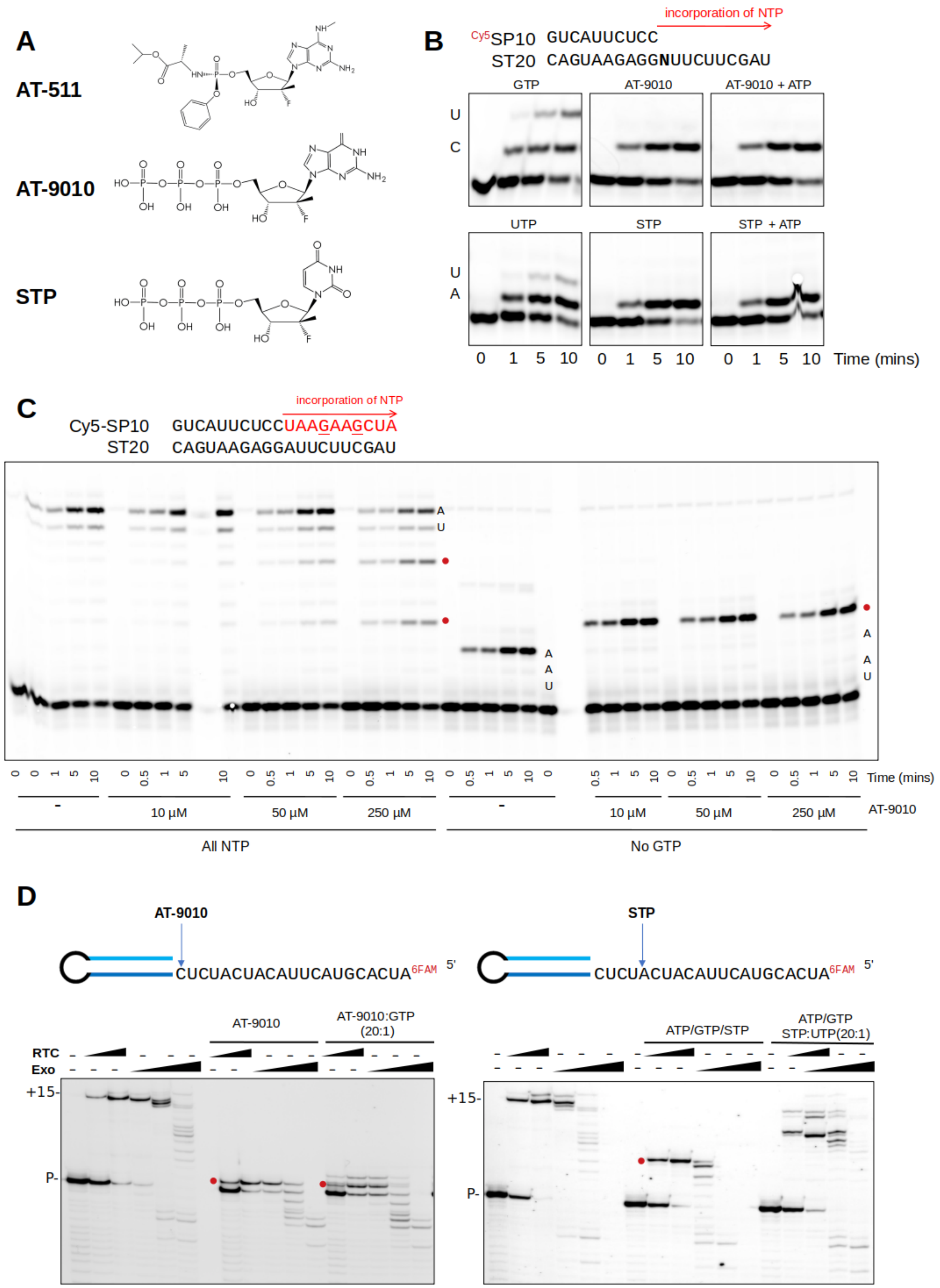
A
AT-511
AT-9010
STP
B
Cy5SP10 GUCAUUCUCC
ST20 CAGUAAGAGGNUUCUUCGAU
incorporation of NTP
GTP
AT-9010
AT-9010 + ATP
UTP
STP
STP + ATP
0 1 5 10 0 1 5 10 0 1 5 10 Time (mins)
C
incorporation of NTP
Cy5-SP10 GUCAUUCUCCUAAGAAGCUA
ST20 CAGUAAGAGGAUUCUUCGAU
Time (mins)
10 µM
50 µM
250 µM
AT-9010
All NTP
No GTP
D
AT-9010
STP
CUCUACUACAUUCAUGCACUA 6FAM 5'
AT-9010:GTP (20:1)
ATP/GTP/STP
ATP/GTP STP:UTP(20:1)
RTC
Exo
+15-
P-

**Figure S5.**

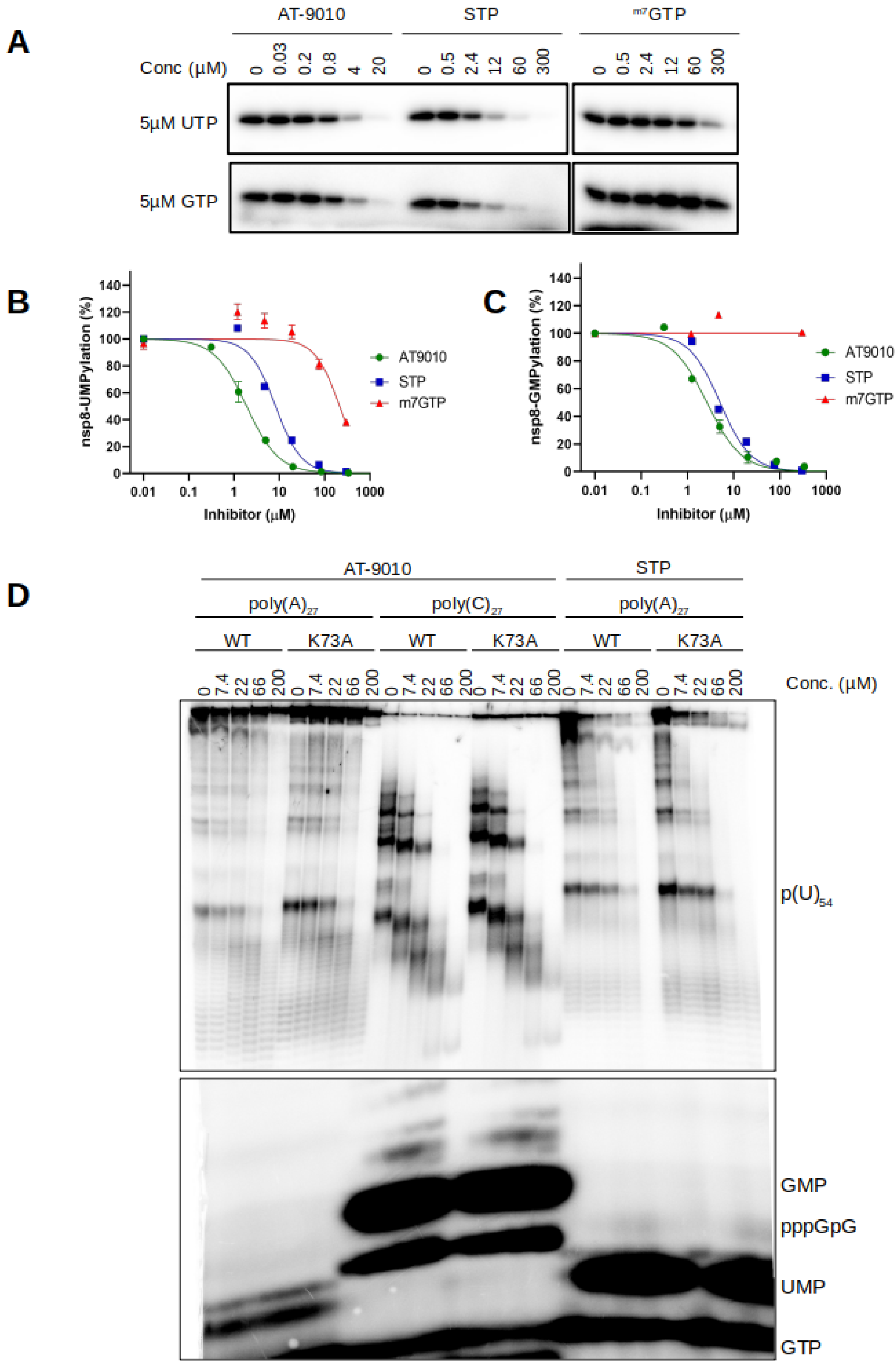
A
AT-9010
STP
m7GTP
Conc (μM)
0 0.03 0.2 0.8 4 20
0 0.5 2.4 12 60 300
0 0.5 2.4 12 60 300
5μM UTP
5μM GTP
B
nsp8-UMPylation (%)
Inhibitor (μM)
AT9010
STP
m7GTP
C
nsp8-GMPylation (%)
Inhibitor (μM)
AT9010
STP
m7GTP
D
AT-9010
STP
poly(A)27
poly(C)27
poly(A)27
WT
K73A
WT
K73A
WT
K73A
0 7.4 22 66 200
Conc. (μM)
p(U)54
GMP
pppGpG
UMP
GTP

**Figure S6.**

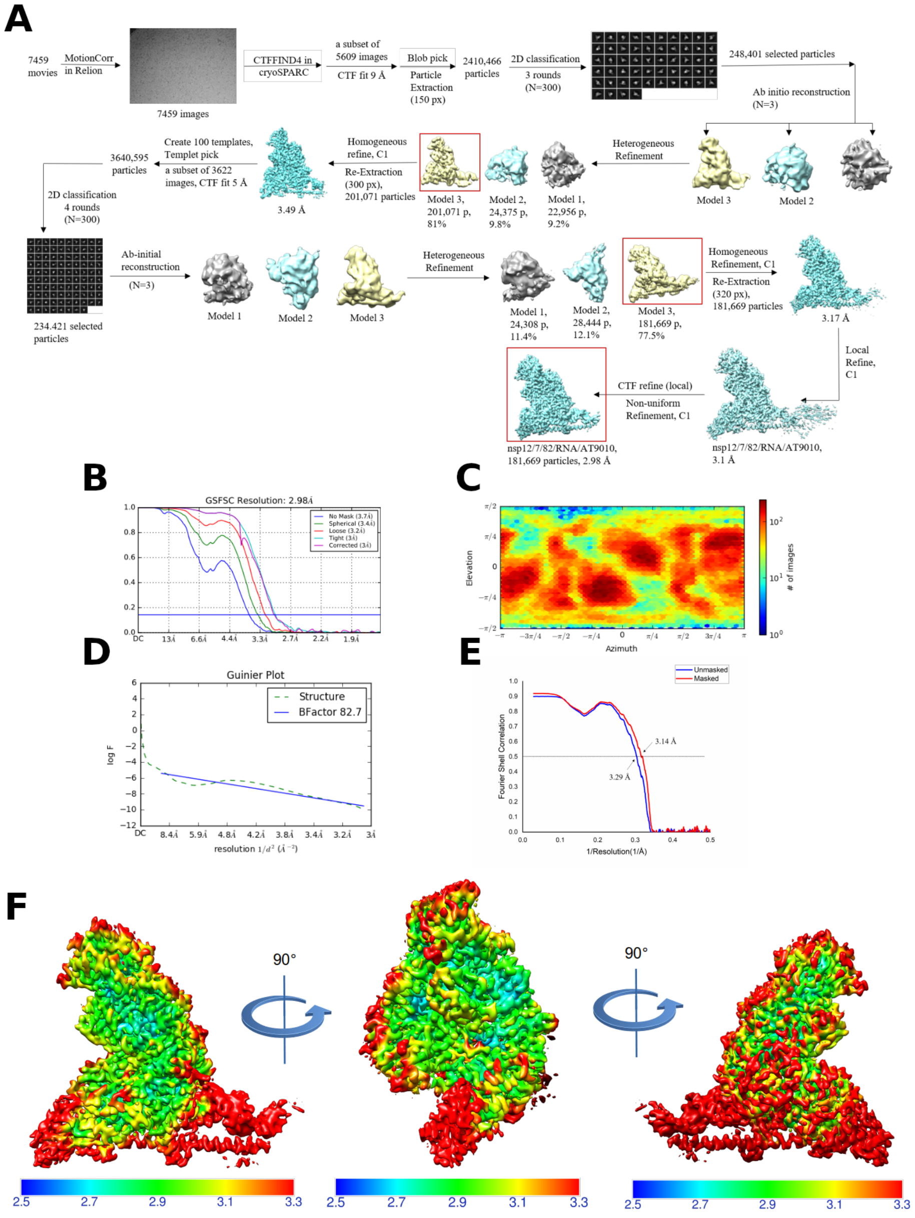
A
7459 movies
MotionCorr in Relion
7459 images
CTFFIND4 in cryoSPARC
a subset of 5609 images
CTF fit 9 Å
Blob pick
Particle Extraction (150 px)
2410,466 particles
2D classification
3 rounds (N=300)
248,401 selected particles
Ab initio reconstruction (N=3)
Model 3
Model 2
Heterogeneous Refinement
Model 3, 201,071 p, 81%
Model 2, 24,375 p, 9.8%
Model 1, 22,956 p, 9.2%
Homogeneous refine, C1
Re-Extraction (300 px), 201,071 particles
3.49 Å
Create 100 templates, Templet pick
a subset of 3622 images, CTF fit 5 Å
3640,595 particles
2D classification 4 rounds (N=300)
234.421 selected particles
Ab-initial reconstruction (N=3)
Model 1
Model 2
Model 3
Heterogeneous Refinement
Model 1, 24,308 p, 11.4%
Model 2, 28,444 p, 12.1%
Model 3, 181,669 p, 77.5%
Homogeneous Refinement, C1
Re-Extraction (320 px), 181,669 particles
3.17 Å
Local Refine, C1
CTF refine (local)
Non-uniform Refinement, C1
nsp12/7/82/RNA/AT9010, 181,669 particles, 2.98 Å
nsp12/7/82/RNA/AT9010, 3.1 Å
B
GSFSC Resolution: 2.98Å
No Mask (3.7Å)
Spherical (3.4Å)
Loose (3.2Å)
Tight (3Å)
Corrected (3Å)
C
Elevation
Azimuth
### of images
D
Guinier Plot
Structure
BFactor 82.7
log F
resolution 1/d² (Å⁻²)
E
Unmasked
Masked
Fourier Shell Correlation
1/Resolution(1/A)
3.14 Å
3.29 Å
F
90°
90°
2.5
2.7
2.9
3.1
3.3

**Figure S7.**

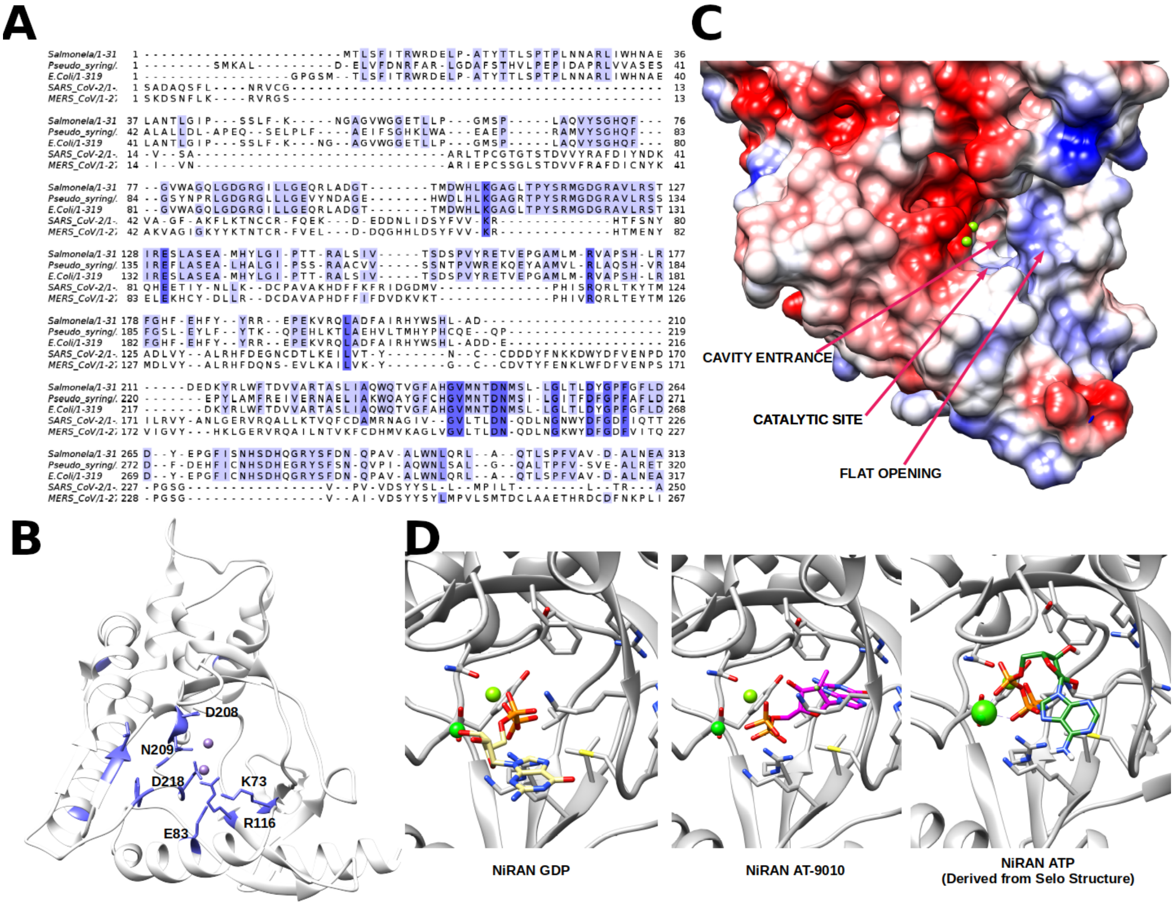
A
B
C
D
CAVITY ENTRANCE
CATALYTIC SITE
FLAT OPENING
D208
N209
D218
K73
E83
R116
NiRAN GDP
NiRAN AT-9010
NiRAN ATP
(Derived from Selo Structure)

**Figure S8.**

Table S1 Cryo-EM data collection, refinement, and validation statistics

| **Nsp12-nsp7-nsp8 complex** | **RNA and AT9010 bound** (PDB ID: 7ED5) (EMDB ID: EM-31061) |
|---|---|
| **Data collection and Processing (for each dataset)** | |
| Microscope | Titan Krios |
| Voltage (keV) | 300 |
| Camera | Gatan K3 Summit |
| Magnification | 105,000 |
| Pixel size at detector (Å/pixel) | 0.83 |
| Total electron exposure ($e^-/Å^2$) | 80.5 |
| Number of frames collected during exposure | 50 |
| Defocus range (µm) | -1.5 ~ -2.5 |
| Phase plate (if used) | N/A |
| - phase shift range (in degrees) | N/A |
| - number of images per phase plate position | N/A |
| Automation software | SerialEM |
| Tilt angle | 0 |
| Energy filter slit width (eV) | 20 |
| Micrographs collected (no.) | 7,459 |
| Micrographs used (no.) | 5,609 |
| Total extracted particles (no.) | 3,640,595 |
| **For each reconstruction:** | |
| Refined particles (no.) | 181,669 |
| Final particles (no.) | 181,669 |
| Point-group or helical symmetry parameters | C1 |
| Resolution (FSC 0.143, Å) | 2.98 |
| Resolution range (local, Å) | 2.7-3.3 |
| Map sharpening B factor ($Å^2$) | 82.7 |
| Map sharpening methods | cryoSPARC v2.15.0 |
| **Model composition** | |
| Protein | 1302 |
| Ligands | 8 |
| RNA | 44 |
| **Model Refinement** | |
| Refinement package | PHENIX-1.19_4085 |
| - real or reciprocal space | real space |
| Model-Map CC | 0.84 |
| Model resolution (Å) | 3.14 |
| FSC threshold | 0.5 |
| B factors ($Å^2$) | |
| Protein residues | 69.09 |
| Ligands | 67.51 |
| RNA | 134.15 |
| R.m.s. deviations from ideal values | |
| Bond lengths (Å) | 0.002 |
| Bond angles (°) | 0.538 |
| **Validation** | |
| MolProbity score | 1.9 |
| CaBLAM outliers | 5.13 |
| Clashscore | 7.89 |
| Poor rotamers (%) | 0.09 |
| C-beta deviations | 0.00 |
| EMRinger score (if better than 4 Å resolution) | 3.32 |
| Ramachandran plot | |
| Favored (%) | 92.5 |
| Outliers (%) | 0.31 |

**Figure S1. Labeling of nsp8 by nsp12 WT and NiRAN mutants, in the presence of $MnCl_2$ and $MgCl_2$.** (A-B) SARS-CoV nsp12 NiRAN-mediated labeling of nsp8 with $\alpha^{32}$P-UTP in the presence of varied concentrations of $MnCl_2$, $MgCl_2$ or both ions. Samples were analyzed on 15% SDS PAGE and stained for total protein (top) and exposed to reveal covalently-bound radioactive UMP (bottom gels). A) Reactions performed with nsp12 and nsp8 in the absence of RNA, b) Reactions performed with nsp12:7L8:8 RTC in the presence of poly$(A)_{27}$ RNA. (C) As in (B) but analyzed on 14% acrylamide urea PAGE gels. (D,E) Labeling of nsp8 with $\alpha^{32}$P-UTP (D) and $\alpha^{32}$P-GTP (E) was performed with the nsp12:7:8 complex with various NiRAN alanine mutants. Labeling reactions were performed for 1h at 37°C. (F) Denaturing SDS-PAGE analysis of SARS-CoV-2 RTC with $\alpha^{32}$P-UTP. Lane 1; without RNA, lane 2; protein complex pre-incubated with UTP + $\alpha^{32}$P-UTP prior to addition of poly$(A)_{27}$ RNA, lane 3; protein complex pre-incubated with poly$(A)_{27}$ RNA prior to addition of UTP + $\alpha^{32}$P-UTP, lanes 4-5; as in lane 2 and 3 but digested with PK
